# Learning consistent subcellular landmarks to quantify changes in multiplexed protein maps

**DOI:** 10.1101/2022.05.07.490900

**Authors:** Hannah Spitzer, Scott Berry, Mark Donoghoe, Lucas Pelkmans, Fabian J. Theis

**Author notes:** Equal contribution (Hannah Spitzer, Scott Berry).

## Abstract

Highly multiplexed quantitative subcellular imaging holds enormous promise for understanding how spatial context shapes the activity of our genome and its products at multiple scales. Yet unbiased analysis of subcellular organisation across experimental conditions remains challenging, because differences in molecular profiles between conditions confound differences in molecular profiles across space. Here, we introduce a deep-learning framework called CAMPA (Conditional Autoencoder for Multiplexed Pixel Analysis), which uses a variational autoencoder conditioned on cellular states and perturbations to learn consistent molecular signatures. Clustering the learned representations into subcellular landmarks allows quantitative comparisons of landmark sizes, shapes, molecular compositions and relative spatial organisation between conditions. By performing high-resolution multiplexed immunofluorescence on human cells, we use CAMPA to reveal how subnuclear organisation changes upon different perturbations of RNA production or processing, and how different membraneless organelles scale with cell size. Furthermore, by integrating information across the cellular and subcellular scales, we uncover new links between the molecular composition of membraneless organelles and bulk RNA synthesis rates of single cells. We anticipate that CAMPA will greatly accelerate the systematic mapping of multiscale atlases of biological organisation to identify the rules by which context shapes physiology and disease.

## Introduction

The wide availability of single-cell -omics techniques has rapidly advanced our understanding of cell biology in health and disease^1,2^. Currently, there is a fast growing range of spatially resolved omics methods that move from the multicellular level to the subcellular level^3–5^. These techniques capture information about abundances of multiple molecular species in single cells across large populations or tissues, at the same time as capturing information about how these molecular species spatially interact. This spatial-context information for large numbers of molecular species simultaneously, from the subcellular scale of molecular complexes and small organelles to the multicellular scale of spatial tissue composition holds enormous promise for understanding biological systems in health and disease.

Cells in different cellular states (e.g. different positions of the cell cycle) or from different perturbation conditions show changes in the relative abundance and subcellular localisation of proteins. To quantify subcellular organisation, unbiased pixel clustering approaches have previously been used^3,4^ to identify subcellular regions with similar molecular profiles within individual conditions^3,4^. However, these approaches weigh all proteins equally in the clustering, which typically results in pixels from different conditions being assigned to different classes even though they may represent the same subcellular structure. As an extreme example, if a perturbation completely eliminates a target protein that forms part of the multiplexed image dataset (Fig. 1a), this perturbation-dependent intensity may be the largest difference in the multiplexed pixel profiles. Direct clustering would then identify a completely independent set of clusters for each condition. While this may be useful for identifying differences between perturbations^4^, it is not appropriate for quantifying changes in the internal organisation of cells because it is difficult to relate subcellular landmarks across conditions in an automated manner – a key aspect of systematically comparing how perturbations affect subcellular protein or organelle atlases. We therefore need approaches that can reveal consistent subcellular landmarks despite condition-dependent changes to the abundance and/or relative localisation of some of the proteins that comprise the multiplexed pixel profiles.

**Figure 1:**
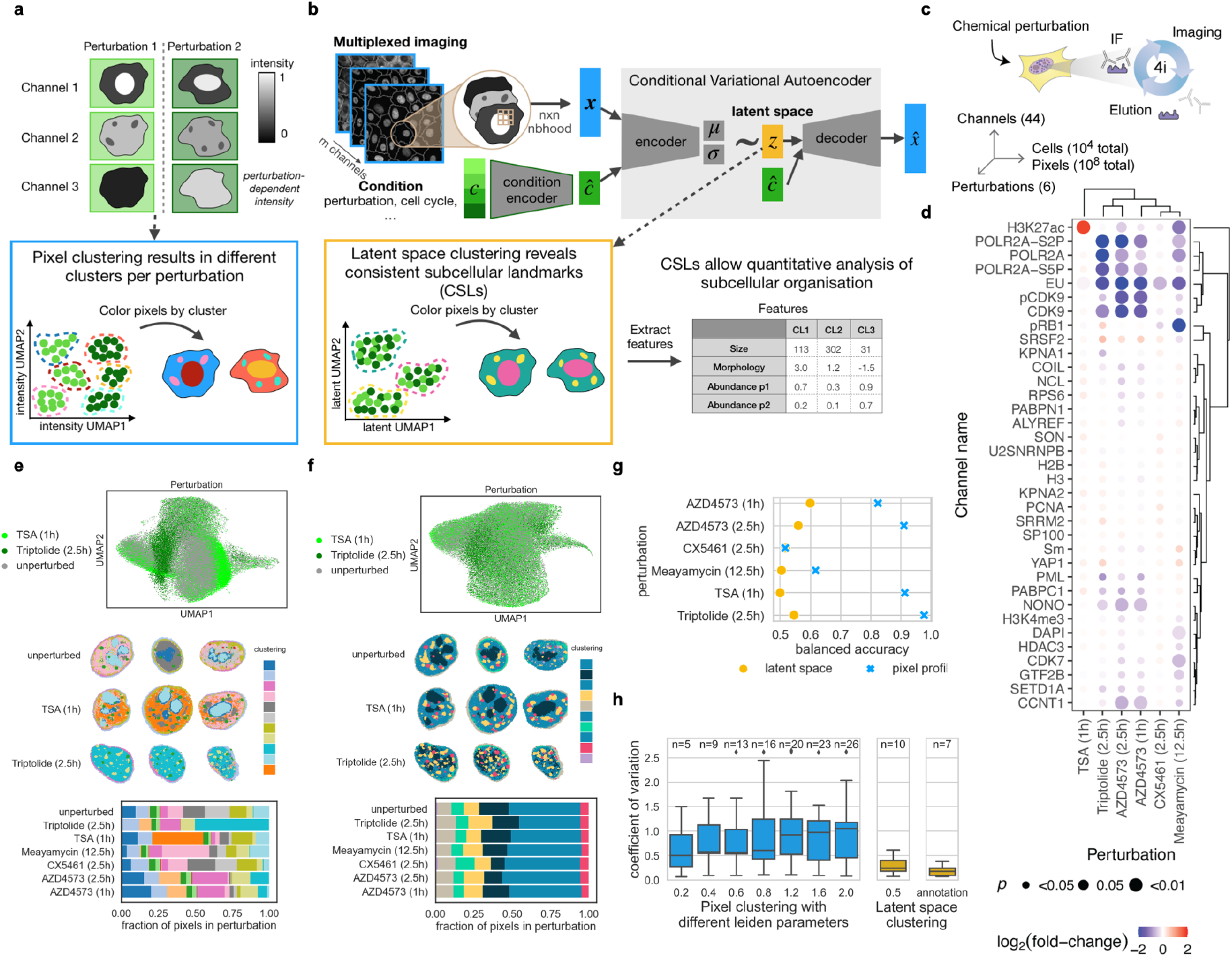
CAMPA enables unsupervised learning of consistent subcellular landmarks using a conditional variational autoencoder. **a:** Schematic of direct pixel clustering across experimental conditions leading to condition-dependent clusters. **b:** Schematic of cVAE conditioned on perturbation to learn a perturbation-independent latent space. Clustering this latent space results in consistent subcellular landmarks (CSLs) from which features can be extracted to qualitatively compare perturbations. **c:** Schematic of 4i experiment and dataset dimensions. **d:** Fold-change in nuclear mean intensity in different perturbations compared to unperturbed cells, for all proteins with nuclear localisation. P-values show significance of perturbation effect on mean intensity, as determined from the mixed effect model (corrected for multiple hypothesis testing using the Benjamini-Yuketeli method). EU indicates ethynyl uridine pulse-labelling of nascent RNA (Methods). **e:** Direct clustering of 4i pixel profiles across perturbations. Top: UMAP representation of multiplexed pixel profiles from unperturbed cells, TSA-treated and Triptolide-treated cells colored by perturbation. Middle: Example cells from each perturbation colored by direct clustering (leiden resolution 1.2). Bottom: Fraction of pixels assigned to each cluster per perturbation colored by direct clustering (leiden resolution 1.2). For a visualisation of all resolutions used in panel h see Supplementary Fig. 3). **f:** cVAE latent space clustering of 4i data across perturbations. Top: UMAP representation of latent space from unperturbed, TSA-treated and Triptolide-treated cells colored by perturbation. Middle: Example cells from each perturbation colored by latent space clustering (leiden resolution 0.5). Bottom: Fraction of pixels assigned to each cluster per perturbation colored by latent space clustering. **g:** Comparison of perturbation-specificity of 4i pixel profiles and cVAE latent space coordinates. Plots show balanced accuracy scores of binary logistic regression classifiers predicting perturbation from normalised 4i pixel profiles or latent representations of pixels. Accuracy values of 0.5 indicate random chance (perturbation information is not present in the data). **h:** Comparison of perturbation-specificity of direct clustering at different leiden resolutions (0.2, 0.4, 0.6, 0.8, 1.2, 1.6, 2.0) with cVAE latent space clustering. Plots show coefficient of variation of the fraction of pixels in each perturbation assigned to each cluster. Boxplot summarises results for all clusters with resulting cluster number n shown above.

To achieve this, we have developed CAMPA (Conditional Autoencoder for Multiplexed Pixel Analysis), a deep learning framework based on conditional variational autoencoders (cVAE)^6^. CAMPA uses a cVAE for unsupervised learning of condition-independent molecular profile representations to identify consistent subcellular landmarks (CSLs), i.e. multimodal pixel clusters that are conserved across conditions. CSLs are subsequently used to quantify spatial re-organisation of CSLs and morphological and molecular changes of CSLs, resulting in a concise comparison of two or more conditions (Fig. 1b). CAMPA is an open-source python package, with strong links to the single cell transcriptomics analysis software scanpy^7^ and its spatial extension squidpy^8^. It allows for high-throughput analysis of high resolution multiplexed imaging datasets with GPU-accelerated assignment of pixels to CSLs.

Here, we use CAMPA to derive a detailed map of subnuclear organisation in different perturbations, directly from high-resolution iterative indirect immunofluorescence imaging (4i)^4^ data measuring proteins and protein states (e.g. phospho-proteins) involved in transcription, chromatin states, mRNA processing and nuclear export, as well as canonical markers of subnuclear organelles. This allows us to uncover and quantify links between inhibition of mRNA splicing or ribosomal RNA production with the morphology, spatial distribution and composition of subnuclear landmarks. We then apply CAMPA to a second 4i dataset comprising the entire cell to study the scaling properties of subcellular membraneless organelles in relation to cell size.

## Results

### Conditional variational autoencoders allow annotation of consistent subcellular landmarks across conditions

In highly multiplexed images, each pixel is represented as a multiplexed pixel profile: a one-dimensional vector containing the intensity of each marker at that spatial location. To identify consistent types of pixels across different experimental conditions even when some of the underlying channels change, we have developed CAMPA. CAMPA first learns a local, condition-independent representation of multiplexed pixel profiles and subsequently clusters the learned representations into consistent subcellular landmarks (CSLs, Fig. 1b) that occur across experimental conditions. To learn a latent representation of each protein observation, a cVAE is trained on a *n*x*n* neighbourhood of the multiplexed pixel profiles, together with a set of condition labels for each pixel to learn a latent representation of each pixel. Pixels are then grouped together by applying the Leiden algorithm^9^ on a knn graph of the learned latent representations. Because the condition labels are supplied as additional input, the cVAE latent representations are mostly condition-independent, and thereby lead to less condition-dependent pixel clusters (cf Fig. 1g,h).

A key aspect of perturbation experiments is to identify and quantify changes in cellular phenotypes that occur upon treatment. Here, we focus on how perturbation of different stages of RNA metabolism affects subcellular organisation, by collecting a high-resolution (pixel size: 108nm x 108nm) 44-plex image dataset of 11,848 human epithelial cells derived from mammary gland (184A1) across six chemical perturbations using 4i^4^ (Fig. 1c). The selected chemical perturbations target different pathways involved in RNA production and processing (histone deacetylation: Trichostatin A (TSA), Pol I transcription: CX5461^10^, Pol II transcription initiation: Triptolide^11^, and activation: AZD4573^12^ and mRNA splicing: Meayamycin^13^). The proteins and post-translational modifications imaged (Supplementary Table 1) either play roles in RNA metabolism, or are molecular markers characteristic of certain cellular states (e.g. cell cycle stage, cell crowding). We observed changes in overall protein state abundances across all perturbations (Fig. 1d), confirming our previous observations in other cell lines^14^. However, we also noticed perturbation-induced changes in the composition and relative spatial organisation of membraneless nuclear organelles involved in RNA metabolism such as nuclear speckles, PML bodies, and the nucleolus. Thus, this dataset provides an ideal use case for the CAMPA framework to generate novel insights into relationships between RNA metabolism and subcellular organisation.

To identify perturbation-dependent changes in composition and spatial organisation of subnuclear membraneless organelles, we initially focused on classifying the approximately 100M nuclear pixels for the 34 markers that localised to the nucleus (Supplementary Fig. 1, Supplementary Tables 2-3). We applied CAMPA cVAE training and clustering on these data using cell-cycle stages (annotated independently of CAMPA^14,15^) and perturbation conditions as categorical condition labels. As expected, we found that multiplexed pixel profiles were highly perturbation-dependent when plotted using a UMAP-embedding^16^ (Fig. 1e). In contrast, when we plotted a UMAP of pixels using their coordinates in cVAE latent space, all conditions appeared to have overlapping distributions. We quantified the condition-independence of the latent representation using binary linear classifiers trained to distinguish pixels from perturbed and unperturbed cells based on their latent representation for all perturbations. These classifiers were often not better than random chance (median accuracy 0.53; min=0.50; max=0.60). In contrast, linear classifiers based on multiplexed pixel profiles reached a median accuracy of 0.87 (min=0.52; max=0.98) (Fig. 1d). We found that using a 3×3 neighbourhood of each pixel as cVAE input, improved cVAE latent space robustness to single-pixel noise which often occurs in microscopy imaging (Supplementary Fig. 2), representing another advantage over direct pixel clustering approaches.

Clustering the latent space (Leiden resolution 0.5) resulted in ten clusters. To accelerate clustering and to enable interactive clustering on a standard workstation, we clustered a subsample of pixels (150k pixels) and then projected resulting clusters to all pixels using 15 nearest neighbours. Using the adjusted mutual information score and adjusted rand index to compare clusterings for different subsamples and leiden initialisations to our final clusters, we found that cluster stability was not significantly influenced by a different random subsample nor by reducing or increasing the number of samples used for the clustering by a factor of two (Supplementary Fig. 2a-b). For comparison with previous approaches, we also directly clustered pixels using their multiplexed pixel profiles^4^. Whereas the intensity space clusters were generally enriched in different perturbations (Fig. 1e), the ten clusters derived from the latent space were evenly distributed across perturbations (Supplementary Fig. 3). To assess the perturbation-specificity of the clusters, we computed the median coefficient of variation (CV) of the fraction of pixels assigned to each cluster across perturbations. The median CV of the latent space clustering is 0.24 (min=0.08; max=0.61), indicating that clusters show similar relative abundance in different perturbations, whereas a direct pixel clustering at similar resolution results in a median CV of 0.57 (min=0.09; max=2.62) (Fig 1h). In addition, despite differences in intensities of some 4i-markers across different cell-cycle phases (e.g. PCNA, pRB1), due to the inclusion of cell-cycle as condition in CAMPA, latent representation contained less cell cycle information (median accuracy of pairwise binary classifiers of latent space/pixel profiles: 0.58/0.67), and latent space clusters were consistently assigned across cell cycle stages (median CV across cell cycle stages of latent space clustering/direct pixel clustering: 0.11/0.21) (Supplementary Fig. 4).

To annotate the latent space clusters, we considered the relative molecular abundance of each 4i-marker in each cluster as well as their spatial distributions within the cell (Fig. 2b-i and Supplementary Fig. 5). In this way, we assigned the ten latent space clusters to seven consistent subcellular landmarks (CSLs), by merging some of the latent clusters into one CSL (Supplementary Fig. 5). The following CSLs were identified: Nucleolus, Nuclear speckles, PML bodies, Cajal bodies, Nucleoplasm, Nuclear periphery, and Extra-nuclear (outside the nucleus) (Fig. 2d-i). To quantitatively validate CSL pixel assignments, we performed two manual segmentations of nuclear speckles and two manual segmentations of PML bodies using state-of-the-art pixel classifiers^17^ (Supplementary Fig. 6). These were based only on single-channel intensities of canonical markers for these membraneless organelles (SON and SRRM2 for nuclear speckles and SP100 and PML for PML-bodies). We quantitatively compared these manual segmentations with their respective CSLs and found that CSL-Nuclear speckles derived were equally similar to the manual segmentations (*F*_1 (*CSL*|*SON*)_ = 0.963 + 0.006, *F*_1(*CSL*|*SRRM*2)_ = 0.967 + 0.006, mean ± s.d. between conditions) as the different manual segmentations are to one another (*F*_1 (*SRRM2*|*SON*)_ = 0.964 + 0.007) (Supplementary Fig. 6). F_1_-scores were similarly high for PML bodies.

**Figure 2:**
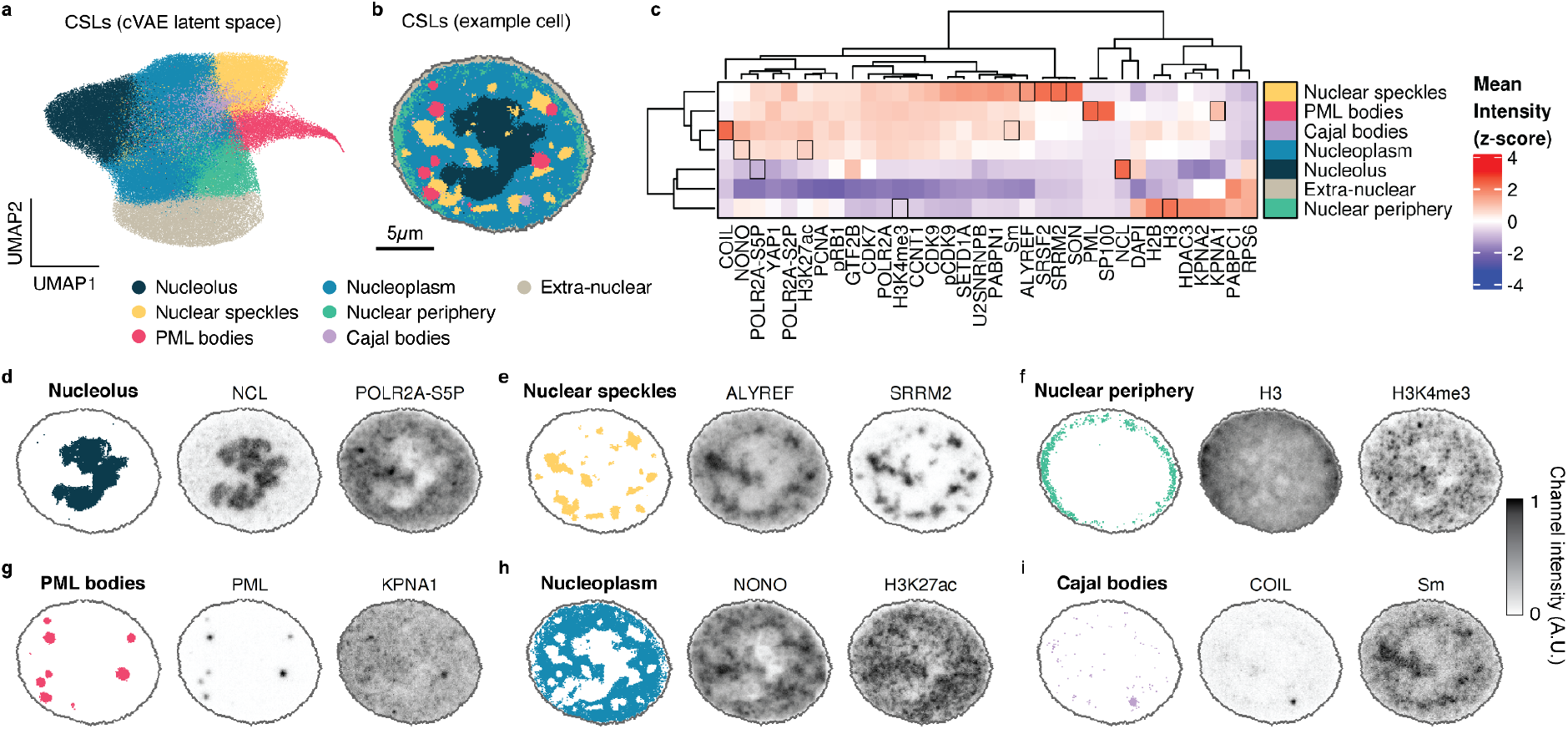
Consistent subcellular landmarks represent known subnuclear structures. **a:** UMAP representation of cVAE latent representations of pixels generated within CAMPA, colored by CSL. **b:** Example cell showing spatial distribution of CSLs. **c:** Relative mean intensity of each channel in each annotated CSL (c.f. Supplementary Fig. 5 for all 10 leiden clusters). Heatmap z-scored by column to show relative localisation of each channel across CSLs. Black boxes highlighted in d. **d-i:** Example 4i channels that are enriched or depleted in the identified CSLs, shown together with CSLs. See c for distribution of channels across CSLs.

We therefore conclude that, in contrast to direct pixel clustering, CAMPA allows unbiased identification and consistent annotation of subcellular landmarks across conditions. Unlike for manual segmentation of subcellular structures, when using CAMPA to identify CSLs, there is no need to pick or pre-define markers of certain landmarks in advance, because the cVAE uses all channels that are consistent across all perturbations to define the latent space. This may ultimately allow identification of novel landmarks defined by higher-dimensional combinations of different channels. Importantly, channels that show characteristic changes in intensity between conditions are ignored by the cVAE when generating the latent space, and are therefore not used to define the subcellular landmarks. Naturally, these channels can then be used to compare the effects of, and differences between perturbations when aggregated on the perturbation-independent landmarks.

### CAMPA uncovers perturbation-induced changes in molecular composition and spatial organisation of subcellular landmarks

To quantify the average subcellular changes in abundance of the proteins and post-translational modifications measured by 4i across the six chemical perturbations, we calculated the mean intensity of each marker in each CSL on a per-cell basis, across all 11,848 cells in the dataset. We then computed the fold-change for a particular condition compared to unperturbed cells across all CSL/channel combinations. We also calculated the fold-change in the size of each CSL, and combined this information into a heatmap-style dot-plot (Fig. 3a, Supplementary Fig. 7a). This generalises the traditional analysis based on nuclear and cell segmentation (Fig. 1d) to additional organelles. Focusing on Meayamycin, which perturbs mRNA splicing^13^, the heatmap predominantly revealed a set of markers that were uniformly depleted across the nucleus, and an overall increase of the relative size of nuclear speckles. To reveal preferential relocalisation of proteins rather than overall changes in abundance, we normalised the fold-changes in each CSL by the whole-nucleus fold-changes (Fig. 3b, Supplementary Fig. 7b). This showed that not only is the relative size of nuclear speckles increased upon Meayamycin treatment, but also their molecular composition changes: they become significantly enriched in cytoplasmic poly(A) binding protein 1 (PABPC1) (Fig. 3e), and depleted in POLR2A-S2P (a marker of actively transcribing RNA Polymerase II) (Fig, 3f). PABPC1 relocalisation to nuclear speckles was observed previously^18^. POLR2A-S2P is normally distributed throughout the nucleoplasm with slight enrichment in nuclear speckles (see Fig. 2b)^19^. However, upon inhibition of mRNA splicing, we find that POLR2A-S2P is reduced in overall abundance (Fig. 3a,f) and, in addition, is specifically excluded from nuclear speckles (Fig. 3b,f). These changes in POLR2A-S2P were mirrored by a reduction in bulk RNA production rate upon Meayamycin treatment, as measured by ethynyl-uridine (EU) pulse labelling (Fig. 1d). Many mRNA splicing factors reside in nuclear speckles and transcription and splicing has been reported to occur more efficiently in their vicinity^20–22^. Moreover, Ser2-phosphorylation of POLR2A is important for coupling of mRNA splicing and transcriptional elongation^23^. However, our analysis shows that the relative abundance of CDK9 (the kinase predominantly responsible for POLR2A-S2P) actually increases within nuclear speckles at the same time (Fig. 2b). This indicates that inhibition of splicing affects both overall transcription rates, and either causes relocalisation of transcribing Pol II (POLR2A-S2P) further from nuclear speckles, or preferentially affects transcription of genes that are normally transcribed in the vicinity of nuclear speckles. This is in agreement with a model in which splicing and transcription are functionally and kinetically coupled^24^.

**Figure 3:**
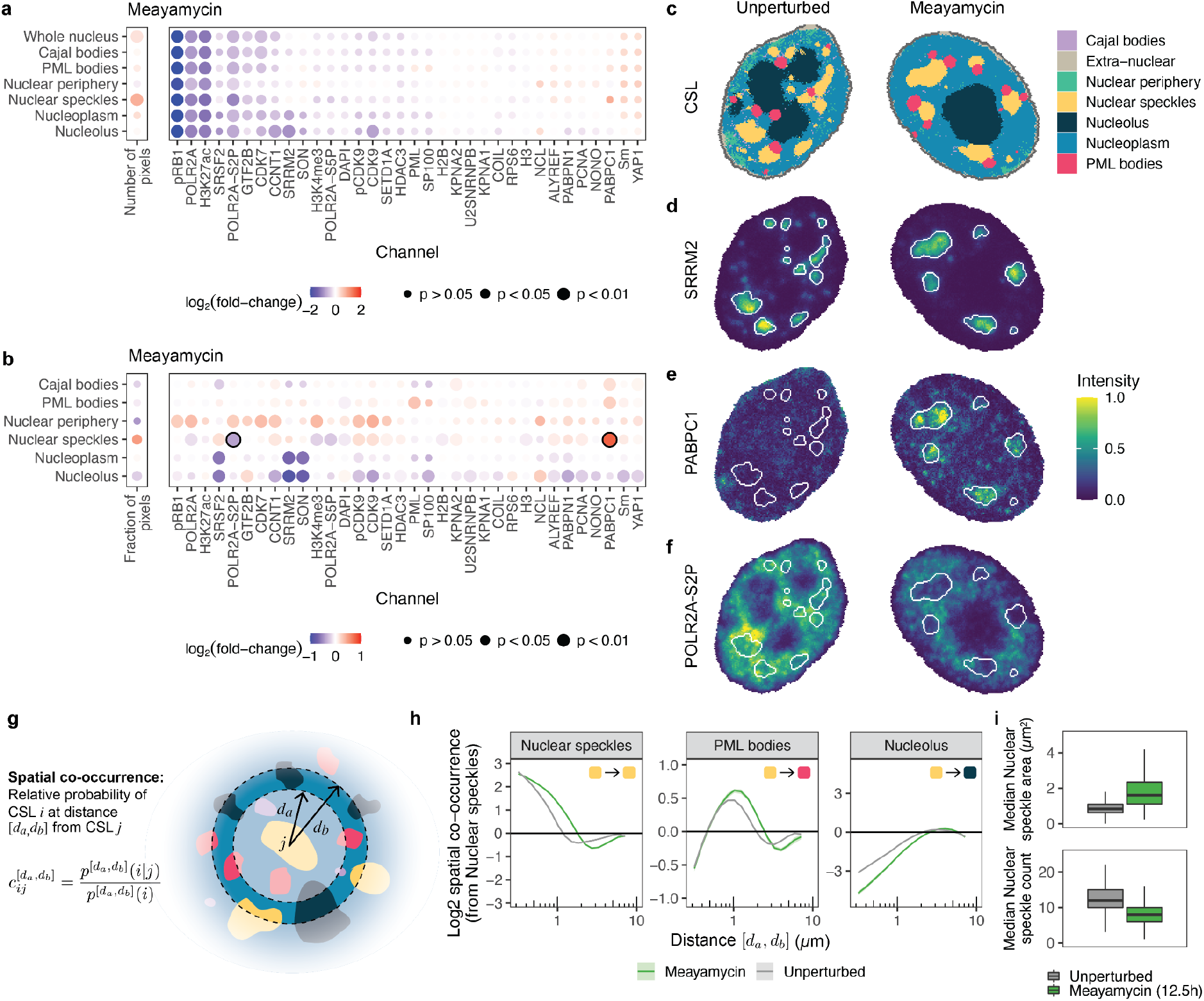
Molecular composition and spatial organisation of subcellular landmarks change upon inhibition of mRNA splicing. **a:** Log2 fold-change of mean intensities for each channel in each CSL, or number of pixels in each CSL, when comparing Meayamycin with unperturbed cells. P-values show significance of Meayamycin treatment on protein levels for each channel/CSL, as determined from the mixed effect model. P-values are corrected for multiple hypothesis testing using the Benjamini-Yuketeli method. **b:** As above, except normalised by the overall (whole-nucleus) changes in intensity. In this case p-values indicate significance of mean intensity change in CSL compared to the change observed for the whole nucleus (see Methods). **c:** Example unperturbed (left) and Meayamycin-treated (right) cell colored by CSL. **d-f:** Example cells from (c) with pixels colored by (d) SRRM2 intensity, (e) PABPC1 intensity, and (f) POLR2A-S2P intensity. **g**: Schematic showing calculation of spatial co-occurrence. **h:** Mean Log2 spatial co-occurrence from Nuclear speckles to Nuclear speckles (auto-co-occurrence); Nucleolus; and PML bodies, as a function of distance on x-axis (on log scale) in Meayamycin-treated and unperturbed cells. Shaded regions indicate 95% confidence intervals for the mean. See Supplementary Fig. 8b for all co-occurrence plots of Meayamycin-treated and unperturbed cells. **i:** Median area and number of individual nuclear speckles per cell. Boxplots summarise distributions over Meayamycin-treated and unperturbed cells. Colours as in h.

In addition to capturing quantitative information about molecular abundances in different subcellular compartments, this dataset has an additional layer of information in the relative spatial arrangement of the identified CSLs. To analyse this, we quantified the spatial organisation within each cell by computing the pairwise spatial co-occurrence between CSLs (Fig. 3g). Spatial co-occurrence^8,25^ captures the relative probability that two CSLs are found within a distance interval from one another (Fig. 3g). Spatial co-occurrence can be averaged over cells in a specific condition (e.g. perturbation) and plotted as a function of distance intervals (Fig. 3h, Supplementary Fig. 8). These plots reveal a fingerprint of how spatial structure changes on average in a condition. At short distances, co-occurrence scores from a structure to itself (auto-co-occurrence scores) are typically high, reflecting the fact that pixels in close spatial proximity are likely to be from the same CSL. For example, comparing Meayamycin-treated cells with unperturbed cells, we found that spatial auto-occurrence of nuclear speckles remains at high values at larger distances in Meayamycin-treated cells. This indicates that the average size of nuclear speckles increases in this perturbation (Fig. 3c), which we confirmed by computing the median area of nuclear speckles per cell (Fig. 3i). Examining the co-occurrence *between* CSLs, we found that co-occurrence of PML bodies and nuclear speckles increases at smaller distances in Meayamycin-treated cells compared to unperturbed cells (Fig. 3h), indicating that PML bodies are more likely to be found in close proximity to nuclear speckles. The opposite effect was observed between nuclear speckles and the nucleolus (Fig. 3h). These spatial relationships were evident in images: upon Meayamycin treatment PML bodies appear to coalesce onto nuclear speckles, and the nucleolus and nuclear speckles appear to move further from one another (Fig. 3c). To our knowledge, neither of these observations have been previously reported. PML bodies have been reported to juxtapose with Cajal bodies^26^ and some PML isoforms (produced through alternative splicing) localise to the nucleolar periphery^27^. All of these compartments, along with nuclear speckles, are thought to form through liquid-liquid phase separation^28^. Relocalisation of PML bodies to contact nuclear speckles could therefore represent surface-wetting between these distinct condensates^29^ (Fig. 3c).

Consistent subcellular landmarks derived by CAMPA can thus be used to analyse and statistically quantify both absolute and relative changes in molecular abundance in different cellular structures and to quantify changes in the size, morphological properties and the high-dimensional subcellular spatial organisation of thousands of individual cells.

### CAMPA-derived cell features allow comparisons of spatial and molecular differences across multiple perturbations

So far we have considered comparisons of each perturbation to unperturbed controls. Because both 4i-based immunofluorescence and CAMPA can be applied in high-throughput to tens of thousands of cells, we can extend these analyses to compare multiple perturbations with one another. To do this, we generated a feature vector for each cell containing the mean abundance of each protein in each CSL. We used this as a representation of the specific subcellular-localised abundance of each quantified protein or protein state. In a similar way, we represented the spatial organisation of the nucleus as a feature vector containing the pairwise spatial co-occurrence scores. To determine how these two different aspects of cellular organisation change across all perturbations, we generated a UMAP representation of cells (Fig. 4a-c), based on each of these vectors separately (CSL UMAP and co-occurrence UMAP). As a baseline, we used the mean nuclear intensities of all measured proteins to represent the intensity information available without subcellular resolution (intensity UMAP, Fig. 4a). For the intensity UMAP, cells from different perturbations occupied different regions of the UMAP in most cases, reflecting the perturbation-dependent abundances of measured proteins. To quantify the differences between perturbations in all three cases, we calculated pairwise silhouette scores using each of the three per-cell representations. Silhouette scores are lower when perturbations are more similar (Figs. 4d-f). Using features of mean nuclear intensity, perturbations targeting Pol II transcription (AZD4573, Triptolide) showed low pairwise silhouette scores, indicating common changes in overall nuclear abundance of the proteins and protein states measured. In almost all cases, we observed that pairwise silhouette scores were higher when considering per-CSL intensities (Fig. 4e) instead of whole-nucleus intensities. This indicates that per-CSL intensities provide a more fine-grained characterisation of the cellular phenotype and are therefore better able to distinguish perturbations. In contrast, we found that spatial co-occurrence scores alone were generally less able to distinguish perturbations than mean nuclear intensities (lower silhouette scores). For example, cells treated with the histone deacetylase (HDAC) inhibitor TSA, were distinct from unperturbed cells when using whole-nucleus intensities but highly similar when using co-occurrence scores (Fig. 4g). This indicates a limited change in spatial organisation of the nucleus upon HDAC-inhibition (for the 4i-markers quantified in our experiment), despite hyperacetylation of histones (see Fig. 1d). One notable exception was the comparison of the RNA Pol I inhibitor CX5461 to unperturbed cells. In this case, spatial information was significantly more informative than molecular abundance information when distinguishing perturbations both at the whole nucleus and CSL-level (Fig. 4g). To pinpoint *how* the spatial organisation of CX5461-treated cells differs from unperturbed cells, we computed the differences between pairwise CSL spatial co-occurrences of CX5461-treated cells and unperturbed controls. This revealed that the major source of difference was due to changes in the relative spatial distribution of the nucleolus CSL compared to itself and other CSLs (Fig. 4h, Supplementary Fig. 9). In particular, the nucleolus showed higher spatial auto-co-occurence at low distances and lower spatial auto-co-occurrence at larger distances indicating that the nucleolus adopts a more compact and spatially coherent conformation in CX5461-treated cells. Moreover, we found that pixels assigned to the nucleolus were more likely to be found close to the nuclear periphery. Examining example images directly, we found that CX5461 treatment results in a circularisation (Fig. 4j) and shrinking (Fig. 4j) of the nucleolus and also fragmentation into smaller regions enriched in the nucleolar marker NCL. Since CX5461 inhibits synthesis of ribosomal RNA (rRNA), changes in the morphology of the nucleolus (the site of rRNA transcription) in CX5461-treated cells are not unexpected. Nonetheless, it shows that our unbiased image-based analysis can rapidly identify that the nucleolus is the primary site of activity of this compound, despite the antibody panel not having a marker for the directly targeted protein (RNA Polymerase I). This points to the exciting future possibility of applying CAMPA in a chemical compound screening format to provide clues to the subcellular locations that are relevant for the activity of a particular molecule.

**Figure 4:**
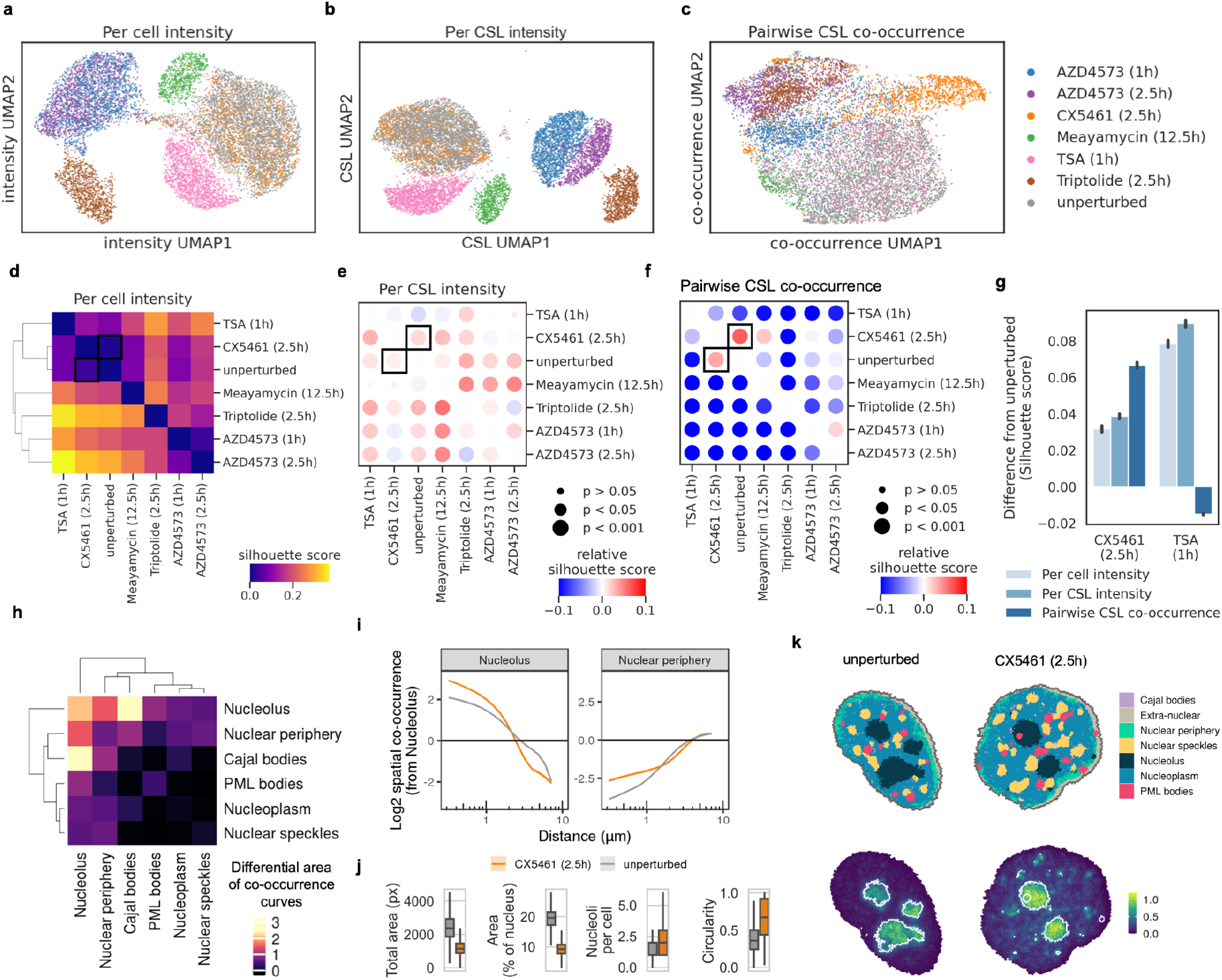
CAMPA-derived cell features allow comparisons of spatial and molecular differences across multiple perturbations. **a-c:** UMAP embedding of cells using different cellular features. Points colored by perturbation. UMAP outliers manually excluded for clarity (Supplementary Fig. 10). Cellular features are (a) per-nucleus mean intensity values, (b) per-CSL mean intensity values, and (c) pairwise CSL spatial co-occurrence scores. **d:** Pairwise differences between perturbations measured by silhouette scores using per-cell mean intensity values. Higher silhouette scores indicate less overlap between perturbations. **e:** Change in silhouette score when considering per-CSL intensities. Negative values indicate decreased silhouette scores compared to per-cell intensity silhouette scores, positive values indicate increased silhouette scores. P-values obtained with Wilcoxon signed-rank test and were adjusted for multiple testing using Bonferroni correction. **f:** As in e, for the change in silhouette score when considering pairwise CSL spatial co-occurrence scores. **g:** Silhouette scores from d-f comparing perturbations with unperturbed controls. Error bars indicate 95% confidence intervals obtained by bootstrapping. **h:** Comparison of pairwise spatial co-occurrences for different CSLs in CX5461 perturbed cells and unperturbed cells quantified as the area between spatial co-occurrences curves. **i:** Mean Log2 spatial co-occurrence from Nucleolus to Nucleolus (auto-co-occurrence) or Nuclear periphery, as a function of distance on x-axis (on log scale) in CX5461-treated and unperturbed cells. Shaded regions indicate 95% confidence intervals for the mean. **j:** Total physical area of Nucleolus in pixels, or as a fraction of the nuclear area, number of nucleoli per cell, median nucleolus circularity per cell. Boxplots summarise distributions across CX5461 and unperturbed cells. Outliers (<Q1 - 1.5xIQR, >Q3 + 1.5xIQR) omitted for clarity. Before obtaining counts and circularity per cell, small objects were removed (Methods). **k:** Top: Example CX5461 and unperturbed cells with pixels colored by CSL. Bottom: Nucleolin (NCL) intensity from the same cells with nucleolus CSL outlines overlaid.

Overall, this analysis reveals that cellular representations obtained through CAMPA can be used to compare cells from several perturbations at once, at the level of subcellular protein localisation or spatial organisation. These are very rich and readily interpretable sources of information, which are complementary to one another and to the overall protein abundance measurements.

### Subcellular reorganisation upon change in cell size

Having developed CAMPA on a 34-plex dataset focused on cell nuclei, we next applied it to whole cell images to demonstrate its potential to identify a larger number of cellular landmarks from higher-dimensional image data. In this case, we examined HeLa cells in which expression of *SBF2* is reduced by siRNA treatment, which results in an approximate 2-fold increase of cell volume and an approximately 3-fold increase in cell area^14^. The average number of pixels per cell in these large cells is 2.1 × 10^5^, compared to 7.8 × 10^4^ pixels for control cells transfected with scrambled siRNA (Supplementary Tables 4-5). We applied CAMPA on 43 channels comprising both nuclear and cytoplasmic 4i stains, using perturbation (SBF2 or scrambled siRNA) and cell cycle stage (G1, S, G2) as conditions (Supplementary Table 6). This resulted in 21 cVAE latent space leiden clusters (Supplementary Fig. 11), some of which were manually merged, resulting in the identification of 16 distinct cytoplasmic and nuclear CSLs (Fig. 5). These CSLs comprised all major compartments marked by the antibodies in the panel, including all previously identified nuclear CSLs as well as cytoplasmic landmarks such as perinuclear and peripheral ER and mitochondria (HSPD1/CALR), Golgi apparatus (GOLGA2) cell-cell contacts (CTNNB1), focal adhesions (PXN) and P-bodies (DDX6) (Fig. 5b-c). Comparing the per-CSL mean intensities of each marker between conditions revealed several differences, however the majority of these were uniform across the whole nucleus or cytoplasm (Supplementary Fig. 12a-b). The more striking changes in this case were differences in the relative sizes of CSLs (Fig. 5d). This indicates that the doubling of cell volume induced by *SBF2* knockdown is associated with disproportionate changes in size of different subcellular compartments. Focusing on membraneless organelles, we found that the markers of the nucleolus and Cajal bodies (NCL and COIL, respectively) both increased their molecular abundance in larger *SBF2*-knockdown cells (Supplementary Fig. 12c-d). However, the size of the nucleolus in *SBF2*-knockdown cells was similar to that of controls (Fig. 5f). Because nuclear area also increases with cell volume upon *SBF2*-knockdown^14^, the size of the nucleolus as a fraction of the nucleus decreases. In contrast, Cajal bodies increased their size by approximately 5-fold - a larger increase than the increase in nuclear or cell area (Fig. 5e). This was predominantly achieved by increasing the size of the individual Cajal bodies, rather than by increasing their number per cell (Fig. 5e, Supplementary Fig. 12-13). In contrast, we found that P-bodies, a cytoplasmic membraneless organelle involved in RNA processing^30^, increased in number per cell rather than by increasing the size of individual P-bodies (Fig. 5g). When we binned cells by cell size, we found that the number of P-bodies in each cell is closely related to cell size, independent of the genetic perturbation (Fig. 5h).

**Figure 5:**
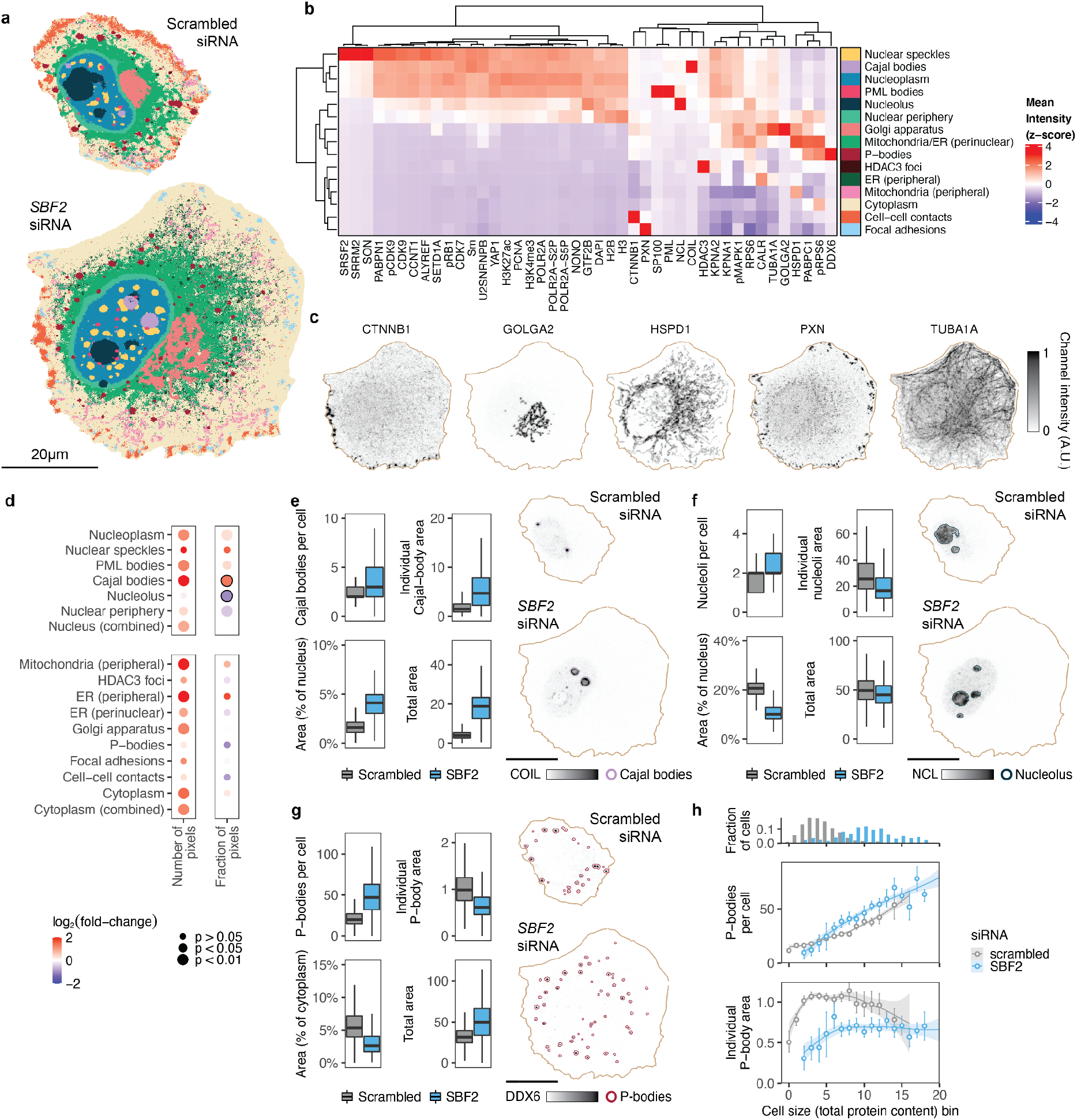
Subcellular landmarks reveal coordination of organelle and cell size. **a:** Consistent subcellular landmarks (CSLs) identified using CAMPA from 43-plex 4i data of HeLa cells transfected with SBF2 siRNA or scrambled siRNA. **b:** Relative mean intensity of each channel in each CSL (cf Supplementary Fig. 11 for all 21 leiden clusters). Heatmap z-scored by column to show relative localisation of each channel across CSLs. **c:** Example 4i images in the example *SBF2*-knockdown cell for comparison with identified CSLs. **d:** Log2 fold-changes of number of pixels per cell assigned to each CSL when comparing *SBF2* knockdown with control cells (scrambled siRNA). P-values show significance of *SBF2* knockdown on abundance of each CSL, as determined from the mixed effect model. P-values are corrected for multiple hypothesis testing using the Benjamini-Yuketeli method. Left panels show non-normalised changes in CSL sizes, right panels show changes normalised to the nuclear or cytoplasmic size-changes, respectively. **e:** Upper: number of Cajal bodies per cell and their per-cell median areas (μm^2^). Before obtaining counts and areas per cell, small objects were removed (Methods). Lower: Cajal body area as a percentage of nuclear area or un-normalised (μm^2^). Boxplots summarise distributions across cells. **f:** As in (e) for NCL / Nucleolus. **g:** As in (e) for DDX6 / P-bodies. **h:** Cells binned by cell size (total protein content). Upper panel shows fraction of cells in each bin per condition. Middle panel shows mean number of P-bodies per cell for each bin. Lower panel shows average size of individual P-bodies (mean of median P-body area per cell). Bins with less than 10 cells per genotype omitted. Error bars show 95% confidence intervals for mean (obtained using bootstrapping). Fit lines show LOESS regression of binned data. Before obtaining counts per cell, small objects were removed (Methods).

This analysis shows that CAMPA generalises to a higher level of multiplexing and is able to identify CSLs not only across conditions with different molecular profiles but also with different CSL sizes. Morphological properties of CSLs on a per-cell basis such as count and area can be used to compare and interpret changes in scaling behaviour between conditions.

### Crossing spatial scales: linking cellular heterogeneity and subcellular reorganisation

Our analysis so far has focused on comparison *between* perturbations, however the analysis of P-bodies as a function of cell size shows that CAMPA can also be used to study how subcellular properties vary *within* populations, potentially allowing the discovery of links between subcellular changes and cellular states.

Rates of RNA production are heterogeneous in cell populations^14,31^ and can be measured by RNA metabolic labelling with 5-ethynyl uridine (EU). Here, we treated cells with a 30 min pulse of EU before fixation and then fluorescently labelled nascent RNA using click chemistry^32^. Nuclear EU intensity quantifies net bulk RNA synthesis rate at the single-cell level (Fig. 6a). To examine how differences in bulk RNA production rates of single cells are related to subcellular changes, we considered control cells (scrambled siRNA) from the CAMPA model trained on entire HeLa cells above (Fig. 5) and binned these into either ‘low’ (lower quartile) and ‘high’(upper quartile) RNA synthesis rates, using mean nuclear EU intensity (Fig. 6b). We then computed the intensity fold-change for each channel/CSL combination between these groups of cells with high and low RNA synthesis rates. This revealed changes in overall nuclear concentration of POLR2A and other proteins related to RNA synthesis (Fig. 6c), as previously observed^14^. Focusing on the subcellular level, we noticed that PML bodies showed a change in relative molecular composition of PML and SP100, the two markers of PML bodies used in this experiment. In cells with low RNA synthesis rates, PML-bodies were enriched in PML, while in cells with high RNA synthesis rates PML bodies were enriched for SP100 (Fig 6c, circled dots). These changes are difficult to observe in overall (all) or whole nucleus (Nucleus (combined)) CSLs, demonstrating the importance of quantifying these changes at the subcellular scale (Fig. 6c). These trends were recapitulated across the full range of EU intensities (Fig 6d), and were observed within G1, S and G2 phases of the cell cycle (Fig. 6e). PML bodies have previously been implicated in transcriptional regulation because they lie near sites of acetylated chromatin and contain transcriptional regulators^33^. However, their molecular composition has not previously been linked to global changes in transcriptional output of single cells. To understand this further we directly examined images of representative cells. This revealed heterogeneity in PML body composition even within a cell. Specifically, cells with low RNA synthesis rates had a subset of PML bodies without SP100, whereas cells with high RNA synthesis rates had a subset of PML bodies without PML. Classically, these bodies are defined as having both SP100 and PML^34^ and are sometimes referred to as ND10 bodies. Univariate annotation of these nuclear bodies based on only PML or SP100 would have not annotated all of these pixels as PML bodies, highlighting a key difference between CAMPA and univariate approaches. It is important to note however that, as we did not use RNA synthesis rate as condition in the cVAE training, we would expect to see these unique combinations of pixels annotated as different CSLs at a higher resolution of cVAE latent space clustering. As expected from the images, a UMAP representation of all pixels from PML bodies using the cVAE latent representation confirmed that there is a spectrum of PML-body pixels ranging from SP100-enriched to PML-enriched that located in different areas of the UMAP space (Supplementary Fig. 15). These results demonstrate that CAMPA can be used not only to reveal changes between perturbations but also to cross the cellular and subcellular scales to uncover links between global properties of cells and subcellular organisation.

**Figure 6:**
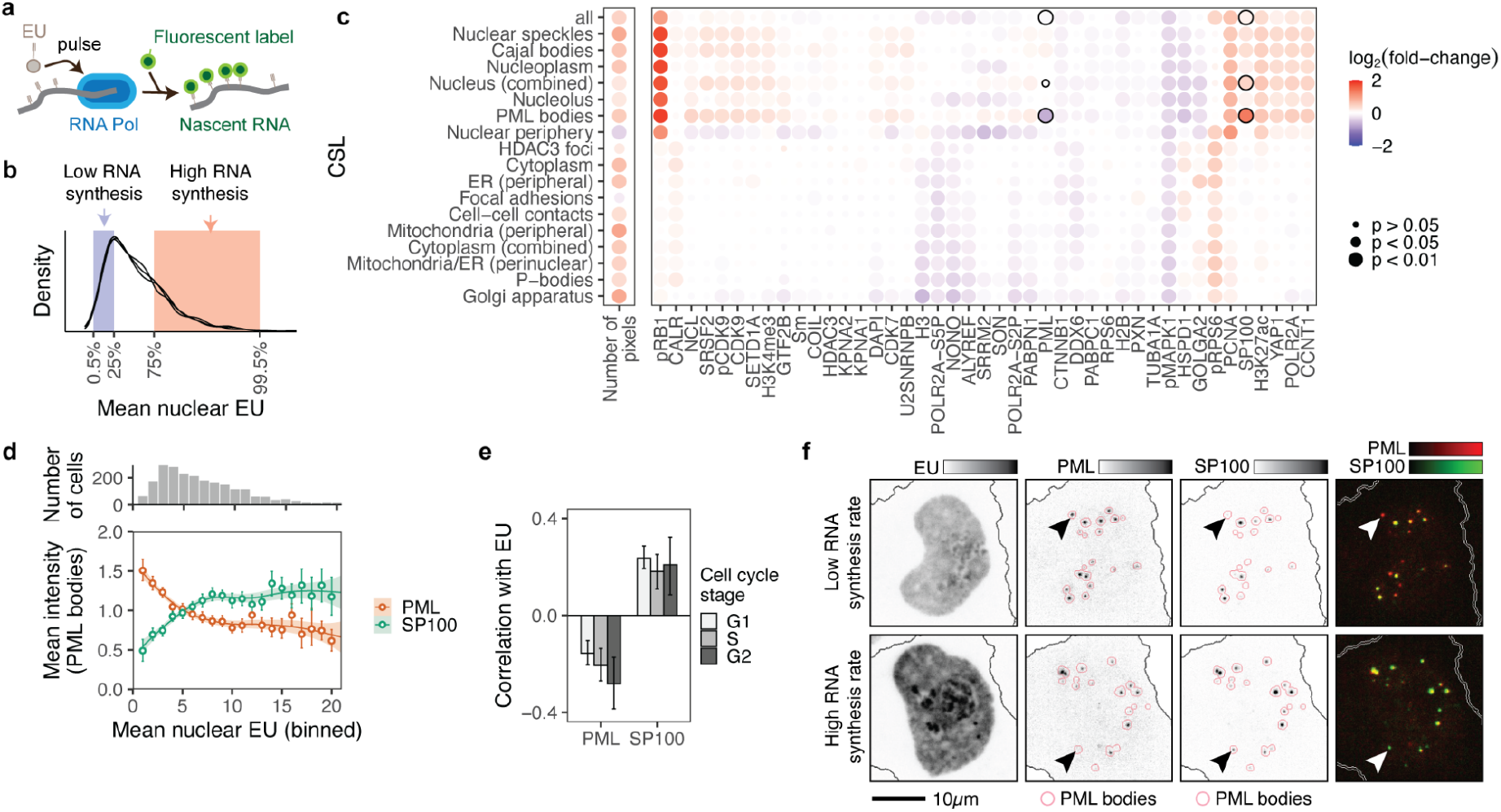
Cellular RNA synthesis rates are associated with altered molecular composition of PML bodies. **a:** Schematic of RNA metabolic pulse labelling with ethynyl uridine (EU)^32^. **b:** Distribution of RNA synthesis rates in the cell population, as quantified by mean nuclear EU incorporation. Cells are binned into upper and lower EU intensity quartiles. **c:** Log2 fold-change of mean intensities for each channel in each CSL, or number of pixels in each CSL, when comparing HeLa cells with high RNA synthesis rates to cells with low RNA synthesis rates. P-values show significance of the difference in mean protein abundance for each channel/CSL, as determined from the mixed effect model. P-values are corrected for multiple hypothesis testing using the Benjamini-Yuketeli method. **d:** Mean intensity of PML and SP100 in PML bodies as a function of RNA synthesis rate (mean nuclear EU intensity) expressed relative to the mean across all cells. All cells binned by EU. Bins with less than 10 cells omitted. Error bars show 95% confidence intervals for mean (obtained using bootstrapping). Fit lines show LOESS regression of binned data. Upper panel shows number of cells in each bin. **e:** Correlations of mean PML-body intensity of PML and SP100 with overall cellular RNA synthesis rate (mean nuclear EU) in each cell cycle phase. Error bars show 95% confidence intervals for correlation (obtained using bootstrapping). **f:** Example images comparing two S-phase cells in states of high and low RNA synthesis. Outlines of PML bodies derived from PML body CSL were dilated by 9 pixels for visualisation purposes.

## Discussion

Quantifying changes in subcellular organisation across perturbations in an automated manner is a central goal in the analysis of high-dimensional multiplexed imaging datasets. This has so far been difficult because perturbation-induced changes or heterogeneity in cell populations has prevented the consistent annotation of subcellular structures in all cells. In the CAMPA framework, we use a cVAE to learn robust perturbation- and cell-state-independent latent representations of pixels which allows the identification of consistent subcellular landmarks (CSLs) that are shared across perturbations and cell states. This differs from previous approaches based on direct clustering of multiplexed pixel profiles, which aim to identify pixel combinations that are unique or enriched in different experimental conditions or cell states and then use these to understand subcellular organisation. In contrast, we use CAMPA to identify consistent types of pixels *across* different conditions and then quantify changes in all markers with respect to these subcellular landmarks. This leads to a more interpretable and quantitative assessment of changes between conditions that directly provides insights into changes in subcellular protein abundance and localisation, and the relative positioning of organisational units in the cell at all subcellular length scales. Compared with direct pixel clustering, CAMPA also scales more readily to comparing large numbers of perturbations, because the number of CSLs that needs to be considered does not necessarily increase with the number of different perturbation conditions studied.

Cellular representations based on CAMPA-derived features can be used to compare multiple perturbations with one another simultaneously. We found that different sources of information (spatial versus intensity-based) were complementary at distinguishing perturbations. Unlike other deep-learning-based approaches for generating cellular representations, the cellular representations we generate are highly interpretable. For example, based on the observation that CX5461-treated cells are distinguishable from unperturbed cells using representations based on subcellular spatial organisation, one can readily identify that this is primarily driven by a change in nucleolar morphology. Because both 4i and CAMPA can be applied in high-throughput, this approach has enormous potential to be applied in screening applications. For example, CAMPA-derived cellular representations could be used as an interpretable fingerprint to characterise and compare perturbations in terms of their subcellular phenotypes.

Here, we focused on subcellular imaging of proteins using 4i, however we anticipate that CAMPA could readily be applied to other modalities such as multiplexed RNA fluorescent in situ hybridisation (FISH)^35^ or integrated spatial genomics^3^ (RNA, proteins and DNA in the same cells) - technologies that have not yet been used to study perturbations at the subcellular scale. However, one limitation of the current CAMPA framework (and all previous pixel clustering approaches) is that pixels are only assigned to one cluster type. The effect of this is that pixel types ‘compete’ for allocation, with markers that show characteristic, sparse distributions in cells preferentially being used to define cellular landmarks. However, limits to optical resolution mean that proteins that do not actually occupy the same physical space in the cell are nonetheless visualised in the same pixels. In our data, the number of structures visualised was appropriate for the optical resolution used, as evidenced by the limited overlap between defining channels of CSLs, however as we further increase the number of structures simultaneously visualised, this problem becomes more pronounced. A possible future extension to CAMPA that may be useful is to use mixture models^36^ or approaches from fuzzy clustering^37^ of the latent space, to allow pixels to be simultaneously assigned to multiple different CSLs.

CAMPA uses a cVAE model to generate consistent latent representations of multiplexed pixel profiles across multiple conditions. Computationally similar approaches have also been used for integrating and clustering cellular representations in single-cell transcriptomics^38,39^. CAMPA’s cVAE model is flexible and could be extended in the future. One example of this would be an adversarial loss to enforce strict disentangling of more complex condition effects and latent representation or ‘architecture surgery’^40^ to allow integration of new data to already learned representations. In this way, CAMPA could contribute to building a queryable atlas of intracellular variation, onto which novel observations from different labs could be projected to not only annotate CSLs, but also to compare with reference atlases. Altogether this will render CAMPA applicable to an even wider range of data and conditions and thus contribute to uncovering the rules by which spatial context shapes the activity of our genome across multiple scales.

## Methods

### Cell lines and culture conditions

HeLa Kyoto (female) cell populations were derived from a single-cell clone and were tested for identity by karyotyping^41^. HeLa cells were cultured in high glucose DMEM supplemented with 10% fetal bovine serum (FBS) and 1% GlutaMAX. Cells with low passage number (2-6) were used for all experiments.

184A1 (human female breast epithelial) cell populations were derived from a single-cell clone, and were used at low passage number (2-6) for all experiments. 184A1 cells were cultured in DMEM/F12 media supplemented with 5% horse serum, 20ng/ml epidermal growth factor, 10μg/ml insulin, 0.5μg/ml hydrocortisone, 10ng/ml cholera toxin.

For all experiments, cells were grown and imaged in uncoated Greiner μClear plastic-bottom 394-well plates.

### Chemical treatments

1250 184A1 cells were plated 72h before chemical treatment. RNA polymerase I inhibitor, CX5461^10^ was dissolved in 5mN HCl at a concentration of 5mM and used at 2μM. XPB (TFIIH) inhibitor, Triptolide^11^ was dissolved in DMSO at a concentration of 10mM and used at 2μM. CDK9 inhibitor AZD4573^12^ was dissolved in DMSO at a concentration of 10mM and used at 0.1μM. Splicing factor 3b subunit 1 (SF3b1) inhibitor, Meayamycin^13^, was dissolved in DMSO at a concentration of 10μM and used at 10nM. Where applicable, final DMSO concentration was 0.1%. Durations of chemical treatments are noted throughout the text and figures.

### siRNA transfection

Transfection with siRNA was performed as previously described^18^. Briefly, 700 HeLa cells were plated per well in 384-well plates for reverse transfection onto a mixture of pooled siRNAs (5 nM final concentration) and Lipofectamine RNAiMAX (0.08μl per well in OptiMEM) according to manufacturer’s specifications. Cells were subsequently grown for 72 hours at 37°C in a final volume of 50μL growth media, to establish efficient knockdown of the targeted genes^14^. *SBF2*-knockdown was validated previously^14^.

### Image acquisition

Imaging was performed an automated spinning-disk microscope (CellVoyager 7000, Yokogawa), equipped with four excitation lasers (405/488/568/647nm) and two Neo sCMOS cameras (Andor), using a 60X/NA1.27 water-immersion objective lens. Band-pass emission filters centred on 445/525/590/675nm were used for detection. Pixel dimensions of images are 108×108nm, with theoretical lateral resolution of 214/252/283/324nm (for emission at 445/525/590/675nm). Images were acquired with a z-spacing of 0.8μm and were maximum-projected during acquisition.

### In situ metabolic labelling of nascent RNA

Cells were pulsed with 5-ethynyl uridine (5-EU) for 30 min before fixation. Nascent RNA was visualised using the Click-iT RNA Alexa Fluor 488 Imaging Kit (Invitrogen), following manufacturer’s instructions except substituting Alexa Fluor 647 azide (Invitrogen) in place of Alexa Fluor 488 azide.

### Iterative indirect immunofluorescence imaging (4i)

4i was performed as previously described^4^ with two modifications: Intercept blocking buffer (LI-COR Biosciences) was used for all blocking, primary and secondary antibody incubations, and 50mM HEPES (Sigma) was included in imaging buffer – which was adjusted to a pH of 7.4. To detect primary antibodies, goat anti-rabbit IgG Alexa Fluor 568 (Thermo Scientific) was combined with either goat anti-mouse IgG Alexa Fluor 488 (Thermo Scientific) or goat anti-rat IgG Alexa Fluor 488 (Thermo Scientific), all at a dilution of 1:500. The first cycle included no primary antibodies, to quantify the background level of fluorescence in all cells. Before 4i experiments, all antibodies were tested for compatibility with elution buffer using the following criteria: similar staining on normal and elution-buffer treated cells, minimal residual signal after elution and re-staining with secondary antibody. The following proteins and post-translational modifications were measured: ALYREF, CALR, CCNT1, CDK7, CDK9, COIL, CTNNB1, DDX6, GOLGA2, GTF2B, H2B, H3, H3K27ac, H3K4me3, HDAC3, HSPD1, KPNA1, KPNA2, NCL, NONO, PABPC1, PABPN1, pCDK9, PCNA, pMAPK1, PML, POLR2A, POLR2A-S2P, POLR2A-S5P, pRB1, pRPS6, PXN, RPS6, SETD1A, Sm antigen, SON, SP100, SRRM2, SRSF2, TUBA1A, U2SNRNPB and YAP1 (Supplementary Fig. 1). Primary antibodies used are listed in Supplementary Table 1.

### DNA and total protein stain

In cycles 1-7, nuclear DNA was stained using 4’,6-diamidino-2-phenylindole dihydrochloride (DAPI) for 5-10 minutes at a final concentration of 0.4μg/mL in phosphate buffered saline (PBS). For cycles 8-22, nuclei were visualised with chicken anti-H2B primary antibody (1:1000, Abcam) and Goat anti-Chicken IgY Alexa Fluor 405 (1:500, Abcam). Before the last imaging cycle, total protein was stained using Alexa Fluor 647 NHS Ester (succinimidyl ester) (Invitrogen) for 10 minutes at a final concentration of 0.2μg/mL in 50mM carbonate-bicarbonate buffer pH 9.2.

### Nuclear and cell segmentation

We typically perform nuclear and cell segmentation as described previously^42^, however this can result in segmentation artefacts when cells are irregularly shaped or highly crowded. To further improve this segmentation, we made use of additional information available in the multiplexed image data. Using DAPI, CALR (endoplasmic reticulum marker) and CTNNB1 (cell-cell contact marker) channels, we manually trained a pixel classifier in Ilastik to identify cell-cell boundaries (which were typically high in CTNNB1 and low in CALR). We refer to the probability map generated as ‘cell outlines’. To segment nuclei, we first used these outlines to mask the DAPI channel and then thresholded and segmented these objects as ‘primary’ nuclei. These were then used as seeds on the original thresholded DAPI image to segment ‘full’ nuclei using propagation. To segment cells, we then summed the total protein and CALR channels and again masked the resulting image with the cell outlines mask to segment ‘primary’ cells. Finally, the primary cells were used as seeds to get the final cell segmentation using a thresholded sum of total protein, CTNNB1 and TUB1A1 channels.

### Data cleanup

After cell segmentation, border cells were excluded. Supervised machine-learning models (support-vector machines) were trained to exclude polynucleated cells and mitotic cells using the TissueMAPS framework^43^, as previously described^14^. After this cleanup we noticed that there were still cells with extreme DNA content. These were removed using manually derived thresholds based on histograms of DNA content. Cells with nuclei that moved during image acquisition, or were incompletely acquired in any cycle were identified and removed by examining the correlation of DNA content at the single-cell level across cycles. The first imaging cycle used a secondary antibody only with no primary antibody. Any cells with excessive background in this staining cycle were also removed from analysis. Supplementary Tables 2 and 4 show the numbers of cells in each of these classes.

### Cell-cycle classification

Cell-cycle classification for 184A1 cells was performed using a machine-learning approach with EdU ground-truth data, as previously described^14,15^. The balanced accuracy of the S-phase classifier was 0.97. For HeLa cells, no EdU wells were included for the SBF2 condition, so no independent ground truth was available. In this case, S-phase cells were manually annotated using PCNA and DAPI texture features by iterative supervised SVM training in the TissueMAPS framework.

### Datasets for cVAE training

Two datasets were collected for training and evaluating cVAE models. Each dataset was split in train, val, and test cells (80%,10%,10% respectively for each dataset). Following the split, multiplexed pixel profiles from the cells were extracted together with their local 3×3 neighbours to make the cVAE latent representation more robust to noise. When one or more of the 3×3 neighbours of the pixel of interest were outside of the segmented region of the cell, the molecular profile of the missing neighbours was replaced with the mean multiplexed pixel profile inside the 3×3 window.

The first dataset consisted of 184A1 cells across six chemical treatments (Supplementary Table 3), using 34 antibody channels localising to the nucleus. For each nucleus in the train and validation split, 0.5% of all molecular profiles were extracted for cVAE training and validation. The second dataset consisted of control and SBF2-knockdown HeLa cells (Supplementary Table 5), using 43 antibody channels, including all those used in the first dataset. For each cell in the train and validation split, 5% of all molecule profiles were extracted for cVAE training and validation. See Supplementary Table 6 for the exact number of cells and molecular profiles in each dataset.

### Preprocessing of datasets

For each channel, background intensity values were calculated using empty wells and subtracted from the molecular profiles. Molecular profiles were normalised using per-channel 98th quantile-normalisation *x_norm_* = *x*/*q*_98_.

This background subtraction and normalisation was also applied to the multiplexed pixel profiles before obtaining a direct clustering.

### Conditional VAE training

The cVAE model consisted of a 3 layer encoder (32,16,16 nodes) with an initial 1×1×32 convolutional layer to mix the channels of individual molecular profile inputs), and a linear layer decoder. Conditions were provided via a 2 layer condition encoder (10, 10 nodes) to the encoder and decoder. The size of the latent representation was 16.

The model was trained to predict the centre pixel of its 3×3 input using σ-VAE^44^: In σ-VAE the variance of the decoder is learned, thus resulting in a calibrated decoder which results in more stable training and better samples from the decoder:

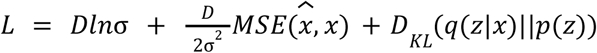

with *x* ∈ *R^D^* the centre pixel of the input, 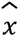 the VAE reconstruction of *x*, and *z* latent space.

The analytical solution for σis given by:

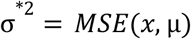

where μ is the estimated latent mean for x.

Training was done for 25 epochs with batch size 128 and learning rate 0.001 (0.0001 for the HeLa dataset). For the 184A1 dataset, the cVAE was trained using perturbation and cell-cycle as conditions by concatenating one-hot encoded representations of both for the condition input. Note that although the control DMSO treatment and untreated cells were used as different conditions in the cVAE model, there was no significant difference in mean intensity between them and they are pooled together for the remainder of the analysis. Together these untreated and DMSO-treated cells are referred to as ‘unperturbed’. For each quantitative comparison between conditions we validated that DMSO and untreated cells showed no differences. These comparisons are shown in Supplementary Figs. 7, 8a and 9. For the HeLa dataset, the cVAE was similarly trained using siRNA condition and cell-cycle as conditions.

### Clustering

For clustering, the dataset was subsampled to 150k (300k for HeLa dataset) multiplexed pixel profiles. To obtain CSLs, a k-nearest neighbour (kNN) graph (k=15) of cVAE latent representations of the subsampled data was computed and partitioned with the Leiden algorithm^9^ using resolution 0.5 (0.9 for HeLa dataset). For comparison, the subsampled multiplexed pixel profiles were also directly clustered by applying Leiden on the kNN graph of the multiplexed pixel profiles with varying resolutions of 0.2,0.4,0.6,0.8,1.2,1.6,2.0. To project cluster assignments to the entire dataset, each data point was assigned to the most frequent cluster within 15 nearest neighbours of the subsampled, clustered set. Neighbours were found using approximate nearest neighbour search^45^.

To assess the impact of subsampling the data before clustering, we varied the random initialisation for the Leiden algorithm (5 different initialisations), the random seed for the subsampling (5 different subsamples), and the size of the subsample, resulting in 5*5 alternative clusterings for each subsample size in 1.1k, 2.3k, 4.6k, 9.3k, 19k, 37k, 75k, 150k, and 300k. The overlap of these clusterings with the final CSLs was computed using the adjusted mutual information (AMI)^46^ and the adjusted rand index (ARI)^47,48^ (cf. Supplementary Fig. 2a-b).

Let *U* = {*U*_1_, *U*_2_,…, *U*_c_} be the groundtruth CSL clustering, and *V* = {*V*_1_, *V*_2_,…, *V*_k_} any other clustering of *n* data points. We calculated AMI(U,V) and ARI(U,V) for all alternative clusterings V to compare clusterings to final CSLs. In addition, we computed the overlap of the resulting clusterings with the final annotated CSLs using the homogeneity score^49^ (cf Supplementary Fig 2c) *h* = 1 – *H*(*U*|*V*)/*H*(*U*), with entropy 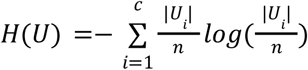.

Homogeneity was calculated for each individual CSL i using a modified 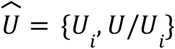 which contained only the CSL i and one other cluster grouping all other CSLs.

To validate CSL pixel assignments, we compared CSLs with manual segmentations of the underlying subcellular structures obtained by training Ilastik^17^ segmentation models on single-channel intensities of canonical markers for these membraneless organelles (cf Supplementary Fig. 6). We quantitatively compared these manual segmentations with their respective CSLs using the F_1_-score (a measurement of classification accuracy) using the manual segmentations as ground-truth 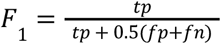, where tp denotes the number of true positives, fp false positives, and fn false negatives.

### Feature extraction using CSLs

For quantitative analysis of differences between conditions, several statistics using the CSLs were computed.

#### Per CSL mean intensity

Per CSL mean intensity values were calculated for each cell and CSL and averaged for each condition.

#### CSL object features

For each cell and CSL, connected components using 8-connectivity were calculated. To filter out noise and get more reliable estimates, only components consisting of more than 10 pixels were counted. In addition, we removed small components from each cell by sorting all components by size and removing the smallest components up to a cumulative area <10% of the total area of the CSL in that cell (Supplementary Figs. 13,14). If no component was smaller than 10% of the total area, no components were removed from that cell.

After filtering, the number, mean or median area, and mean or median circularity of these components was extracted and median-averaged across cells for each condition.

Circularity c was computed as *c* = 4πa/*p*^2^ where a is the area and p the perimeter of the component.

#### Spatial co-occurrence

Spatial co-occurrence^8,25^ *c_ij_*^[*d_a_,d_b_*]^ captures the relative probability that two CSLs (*i, j*) are found within a distance interval [*d_a_, d_b_*] from one another:

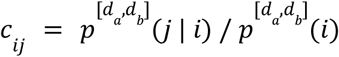

Distance intervals were log-spaced to allow focussing on small-scale changes in spatial reorganisation. For the 184A1 dataset, 19 log-spaced distance intervals between 2 and 80 were used. For the HeLa dataset, 27 log-spaced distance intervals between 2 and 320 were used. The maximum distance of 80 pixels (320 pixels) was chosen to be approximately the 99th quantile of the maximum radius of the nucleus (of the cell for the HeLa dataset).

#### Statistical analysis of mean intensity and CSL abundance changes

To quantify the changes in channel intensities in CSLs, we estimated the fold-difference of each channel between treated and unperturbed control cells in the geometric mean of the per-CSL mean intensity. Specifically, if *Y_ijk_* denotes the mean intensity for CSL *k* within cell *j* of well *i,* we fit a hierarchical linear mixed effects model:

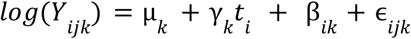

where μ_*k*_ denotes the (log) geometric mean of CSL *k* in the control group and *t_i_* is an indicator variable for condition (*t_i_* = 1 for treated wells and 0 for unperturbed control wells), such that *exp*(*γ_k_*) is the treatment effect on CSL *k*. To account for clustering, β_*i*_~*N*(0, Σ_w_) is a multivariate normal well-specific random effect, with mean zero and general covariance matrix Σ_w_, and ϵ_*ij*_ is a multivariate normal random error with mean zero and covariance matrix Σ_ϵ(*i*)_ = *S_t_i__ RS_t_i__* where *S_t_i__* is a diagonal matrix of (condition-specific) standard deviations of the CSL-specific errors, and *R* is an unstructured correlation matrix that captures the relationships between CSLs within a single cell. Before calculating fold differences and hypothesis testing, we removed compartments of size 0.

For each CSL, we tested the null hypothesis of no treatment effect (γ_k_ = 0) using a Wald test. To reveal relative relocalisation of proteins rather than overall changes in abundance, the fold-changes in each CSL were normalised by the whole-nucleus fold-changes. That is, if *k* = 0 denotes mean intensity across the whole nucleus, then *exp*(γ_*k*_ – γ_0_) is the compartment-specific treatment effect for CSL *k*, and we similarly tested γ_*k*_ = γ_0_ using a Wald test. CSL sizes were analysed in the same way as mean channel intensities.

We used the nlme package^50^ in R version 3.6.3^51^ to fit these models, and used emmeans^52^ to extract estimates and perform the hypothesis tests of interest. For computational efficiency, we fit a separate model for each CSL for each marker (using only the data from that CSL and the whole nucleus), and used the conservative “containment” method^53^ to determine the degrees of freedom of the Wald statistic in the analyses of CSL vs whole-nucleus differences. The false discovery rate was controlled across all combinations of CSLs and channels for each treatment using the Benjamini–Yekutieli method^54^.

### Comparing perturbations

To compare perturbations with respect to different aspects of cellular organisation, we generated three separate cellular representations:

- Mean nuclear intensities of all proteins
- Per CSL mean intensities of all proteins
- Pairwise spatial co-occurrence between CSLs

To measure how “mixed” perturbations are when using these different cellular representations, we calculated silhouette scores^55^ (using L1 distance) S(p,q) for each pair of perturbations p, q:

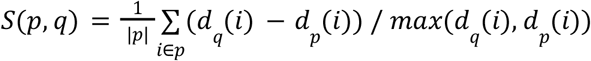

With *d_p_* (*i*) the mean L1 distance of i to all all elements in perturbation p:

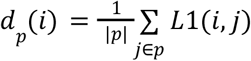

## Code availability

CAMPA is available at https://github.com/theislab/campa with docs at https://campa.readthedocs.io.

All files and scripts necessary for reproducing the results and figures presented in this paper can be found at https://github.com/theislab/campa_ana.

Nuclear and cell segmentation, identification of border cells, and cell-cycle classification was performed using TissueMAPS, an open-source project for high-throughput image analysis which is available at https://github.com/pelkmanslab/TissueMAPS. The TissueMaps analysis pipeline description with module files containing parameter settings used for the preprocessing of data in this paper is provided at https://github.com/theislab/campa_ana.

## Data availability

CSL-derived features from the 184A1 and the HeLa datasets are available at https://doi.org/10.6084/m9.figshare.19699651.

The entire dataset and pre-trained models (~600GB in total) are available upon request.

## Acknowledgements

We thank all members of Lucas Pelkmans’ and Fabian Theis’ lab for discussions and manuscript comments. We thank David Spector for the anti-SNRPB2 and Sm antibodies; Archa Fox for the NONO antibody; Karsten Weis for the KPNA1 and KPNA2 antibodies; and Kazunori Koide for providing Meayamycin. S.B. is supported by an EMBO long-term fellowship (ALTF1175-2016), an HFSP long-term fellowship (LT000238/2017-L) and the University of New South Wales. L.P. is supported by the European Research Council (ERC-2019-AdG-885579) and the Swiss National Science Foundation (SNSF grant 310030_192622). S.B. and L.P. are supported by the University of Zurich. H.S. and F.J.T. acknowledge support by the BMBF (grant# 031L0210A) and by the Helmholtz Association’s Initiative and Networking Fund through Helmholtz AI [grant number: ZT-I-PF-5-01].

F.J.T consults for Immunai Inc., Singularity Bio B.V., CytoReason Ltd, and Omniscope Ltd, and has ownership interest in Dermagnostix GmbH and Cellarity.

## Author contributions

H.S., S.B., L.P., F.J.T. conceived the study and wrote the manuscript; H.S. wrote the CAMPA code; S.B. performed experiments; H.S. and S.B. designed and performed the data analysis; M.D. designed and implemented statistical analysis of mean intensity and CSL-abundance changes; L.P. and F.J.T. supervised the work; all authors read and corrected the final manuscript.

