## Supplements for "Learning consistent subcellular landmarks to quantify changes in multiplexed protein maps"

**Supplementary information**

**Supplementary Table 1: Antibodies and other reagents**

| REAGENT | SOURCE | IDENTIFIER |
|---|---|---|
| Antibodies | | |
| Mouse monoclonal anti-ALYREF | Santa Cruz Biotechnology | Cat#sc-32311; Clone#11G5; Lot#G1517; RRID: AB_626667 |
| Rabbit polyclonal anti-CALR | Abcam | Cat#ab2907; Lot#GR3206421-2; RRID: AB_303402 |
| Mouse monoclonal anti-CCNT1 | Santa Cruz Biotechnology | Cat#sc-271348; Clone#E-3; Lot#J2617; RRID: AB_10608086 |
| Mouse monoclonal anti-CDK7 | Santa Cruz Biotechnology | Cat#sc-7344; Clone#C-4; Lot#L011; RRID: AB_627243 |
| Rabbit monoclonal anti-CDK9 | Cell Signaling Technology | Cat#2316; Clone#C12F7; Lot#7; RRID: AB_2291505 |
| Mouse monoclonal anti-COIL | Abcam | Cat#ab87913; Clone#IH10; Lot#GR50257-2; RRID: AB_10860831 |
| Mouse monoclonal anti-CTNNB1 | Cell Signaling Technology | Cat#2677; Clone#L54E2; Lot#3; RRID: AB_1030943 |
| Rabbit polyclonal anti-DDX6 | Bethyl Laboratories | Cat#A300-461A; RRID: AB_2277216 |

| | | |
|---|---|---|
| Mouse monoclonal anti-GOLGA2 (GM130) | BD Biosciences | Cat#610823; Lot#7163670; RRID:AB_398142 |
| Rabbit polyclonal anti-GTF2B | Atlas Antibodies | Cat#HPA061626; Lot#R86237; RRID:AB_2684570 |
| Chicken polyclonal anti-H2B | Abcam | Cat#ab134211; RRID: N/A |
| Rabbit polyclonal anti-H3 | Abcam | Cat#ab1791; Lot#GR3190126-1; RRID: AB_302613 |
| Rabbit polyclonal anti-H3K27ac | Abcam | Cat#ab4729; Lot#GR3231937-1; RRID: AB_2118291 |
| Rabbit polyclonal anti-H3K4me3 | Abcam | Cat#ab8580; Lot#GR3190162-1; RRID: AB_306649 |
| Rabbit polyclonal anti-H3K9me3 | Abcam | Cat#ab8898; RRID: AB_306848 |
| Rabbit polyclonal anti-HDAC3 | Abcam | Cat#ab7030; Lot#GR166225-20; RRID: AB_305708 |
| Mouse monoclonal anti-HSPD1 (HSP60) | Abcam | Cat#ab13532; Clone#Mab11-13; Lot#GR197160-6; RRID: AB_300433 |
| Rabbit polyclonal anti-KPNA1 | Laboratory of Karsten Weis | RRID: N/A |
| Rabbit polyclonal anti-KPNA2 | Laboratory of Karsten Weis | RRID: N/A |
| Rabbit polyclonal anti-NCL (C23) | Santa Cruz Biotechnology | Cat#sc-13057; Lot#G3114; RRID: AB_2229696 |
| Mouse monoclonal anti-NONO | Laboratory of Archa Fox | RRID: N/A |
| Mouse monoclonal anti-PABPC1 | Santa Cruz Biotechnology | Cat#sc-32318; Clone#10E10; Lot#D1913; RRID: AB_628097 |
| Rabbit monoclonal anti-PABPN1 | Abcam | Cat#ab75855; Clone#EP3000Y; Lot#GR3202568-6; RRID: AB_1310538 |
| Rabbit polyclonal anti-pCDK9 (pThr186) | Cell Signaling Technology | Cat#2549; Lot#2; RRID: AB_2077300 |
| Rabbit monoclonal anti-PCNA | Cell Signaling Technology | Cat#13110; Clone#D3H8P; Lot#4; RRID: AB_2636979 |

| | | |
|---|---|---|
| Rabbit monoclonal anti-pMAPK1 (ERK-pThr202/204) | Cell Signaling Technology | Cat#4370; Clone# D13.14.4E; Lot#24; RRID: AB_2315112 |
| Mouse monoclonal anti-PML | Santa Cruz Biotechnology | Cat#sc-966; Clone#PG-M3; Lot#K1215; RRID: AB_628162 |
| Mouse monoclonal anti-POLR2A (RPB1) | Santa Cruz Biotechnology | Cat#sc-55492; Clone#F-12; Lot#C0119; RRID: AB_630203 |
| Rat monoclonal anti-POLR2A-S2P | Millipore | Cat#04-1571; Clone#3E10; Lot#3169853; RRID: AB_10627998 |
| Rat monoclonal anti-POLR2A-S5P | Millipore | Cat#04-1572; Clone#3E8; RRID:AB_10615822 |
| Rabbit monoclonal anti-pRB1 (pSer807/811) | Cell Signaling Technology | Cat#8516; Clone#D20B12; Lot#8; RRID: AB_11178658 |
| Rabbit monoclonal anti-pRPS6 (pSer235/236) | Cell Signaling Technology | Cat#4858; Clone#D57.2.2E; Lot#16; RRID: AB_916156 |
| Mouse monoclonal anti-PXN | BD Biosciences | Cat# 610052; RRID: AB_397464 |
| Mouse monoclonal anti-RPS6 | Cell Signaling Technology | Cat#2317; Clone#54D2; Lot#4; RRID: AB_2238583 |
| Rabbit polyclonal anti-SETD1A | Atlas Antibodies | Cat#HPA020646; Lot#A96712; RRID:AB_1856752 |
| Mouse monoclonal anti-Sm | David Spector {Pettersson.1984} | Clone#Y12; RRID: AB_2692320 |
| Rabbit polyclonal anti-SON | Atlas Antibodies | Cat#HPA023535; RRID: AB_1857362 |
| Rabbit polyclonal anti-SP100 | Atlas Antibodies | Cat#HPA017384; Lot#A106767; RRID: AB_1857399 |
| Rabbit polyclonal anti-SRRM2 | Atlas Antibodies | Cat#HPA041411; Lot#A90041; RRID: AB_10796671 |
| Mouse monoclonal anti-SRSF2 | Sigma-Aldrich | Cat#S4045; RRID:AB_477511 |
| Mouse monoclonal anti-TUBA1A | Abcam | Cat#ab7291; Clone#DM1A; Lot#GR50257-2; RRID: AB_2241126 |
| Mouse monoclonal anti-U2SNRNPB (B'') | Laboratory of David Spector {Habets.1985} | Clone#4G3; RRID: N/A |

| | | |
|---|---|---|
| Mouse monoclonal anti-YAP1 | Santa Cruz Biotechnology | Cat#sc-101199; Clone#63.7; Lot#K0513; RRID:AB_1131430 |
| Goat polyclonal anti-Mouse IgG (H&L), Alexa Fluor 488 | Thermo-Fisher | Cat#A11029; RRID: AB_138404 |
| Goat polyclonal anti-Rabbit IgG (H&L), Alexa Fluor 568 | Thermo-Fisher | Cat#A11036; RRID:AB_10563566 |
| Goat polyclonal anti-Chicken IgY (H&L), Alexa Fluor 405 | Abcam | Cat#ab175674; RRID N/A |
| Goat polyclonal anti-Rabbit IgG (H&L), Alexa Fluor 488 | Thermo-Fisher | Cat#A11034; RRID: AB_2576217 |
| Goat polyclonal anti-Mouse IgG (H&L), Alexa Fluor 568 | Thermo-Fisher | Cat#A11031; RRID: AB_144696 |
| Goat polyclonal anti-Rat IgG (H&L), Alexa Fluor 568 | Thermo-Fisher | Cat#A11077; RRID: AB_141874 |
| Chemicals, Peptides, and Recombinant Proteins | | |
| Alexa Fluor 647 Azide | Thermo-Fisher | Cat#A10277 |
| Alexa Fluor 647 NHS Ester | Thermo-Fisher | Cat#A20006 |
| Sodium ascorbate | Sigma-Aldrich | Cat#A7631 |
| Copper sulfate | Sigma-Aldrich | Cat#7758-98-7 |
| 5-ethynyl uridine (5-EU) | Baseclick Gmbh | Cat#BCN-003 |
| DAPI (4',6-diamidino-2-phenylindole, dihydrochloride) | Thermo-Fisher | Cat#D1306 |
| Meayamycin | [13] | N/A |
| Triptolide | Adipogen Life Sciences | Cat#AG-CN2-0448 |
| AZD4573 | Selleckchem | Cat#S8719 |
| CX5461 | Axon Medchem | Cat#2173 |

| | | |
|---|---|---|
| Trichostatin A (TSA) | Selleckchem | Cat#S1045 |
| Epidermal growth factor | Sigma-Aldrich | Cat#01-107 |
| Insulin | Sigma-Aldrich | Cat#I1882 |
| Hydrocortisone | Sigma-Aldrich | Cat#H0888 |
| Cholera toxin | Sigma-Aldrich | Cat#C8052 |
| Fibronectin | Sigma-Aldrich | Cat#F0895 |
| Fetal bovine serum | Sigma-Aldrich | Cat#F7524 |
| Horse serum | Gibco | Cat#16050122 |
| DMEM, high glucose | GIbco | Cat#41965062 |
| DMEM/F12 | Gibco | Cat#11330032 |
| OptiMEM Reduced Serum Medium | Gibco | Cat# 31985070 |
| Penicillin/Streptomycin | Gibco | Cat#15140122 |
| Lipofectamine RNAimax transfection reagent | Invitrogen | Cat#13778150 |
| 16% Paraformaldehyde | Electron Microscopy Sciences | Cat#EMS-15710 |
| Intercept blocking buffer | Li-Cor | Cat#927-70001 |
| Triton X-100 | Sigma-Aldrich | Cat#T8787 |
| Experimental Models: Cell Lines | | |
| Human: HeLa, cervical cancer cell line (single-cell clone) | [41] | Kyoto |

| | | |
|---|---|---|
| Human: 184A1, breast epithelial cell line (single-cell clone) | This paper | ATCC CRL-8798 |
| Oligonucleotides | | |
| Negative control siRNA #1; Ambion Silencer Select | Thermo-Fisher | Cat#4390843 |
| siRNAs targeting SBF2; Ambion Silencer Select | Thermo-Fisher | Cat#s37818, s37819, s37820 |
| Other | | |
| 384-well µClear plates | Greiner Bio-One | Cat#781092 |

**Supplementary Table 2: Number 184A1 cells removed during data cleanup across all conditions**

| | n | % total |
|---|---|---|
| **All (non-border) cells imaged** | **13053** | **100%** |
| Mitotic cells (excluded) | 636 | 4.9% |
| Polynucleated cells (excluded) | 343 | 2.6% |
| Extreme DNA content (excluded) | 92 | 0.7% |
| Acquisition error (excluded) | 103 | 0.8% |
| Background artefacts (excluded) | 31 | 0.2% |
| **Final analysed cells** | **11848** | **90.8%** |

**Supplementary Table 3: Number of 184A1 cells and pixels analysed per conditi*on***

| Condition | Number of cells | Number of pixels (nuclei) | Mean area (nuclei) |
|---|---|---|---|
| AZD4573 (2.5h) | 1186 | 1.7E+07 | 1.4E+04 |
| AZD4573 (1h) | 1942 | 2.6E+07 | 1.4E+04 |
| CX5461 (2.5h) | 1552 | 2.0E+07 | 1.3E+04 |
| DMSO (2.5h) | 528 | 6.7E+06 | 1.3E+04 |
| DMSO (12h) | 606 | 7.7E+06 | 1.3E+04 |
| Meayamycin (12h) | 755 | 1.2E+07 | 1.6E+04 |
| Untreated | 2546 | 3.3E+07 | 1.3E+04 |
| Triptolide (2.5h) | 1018 | 1.3E+07 | 1.2E+04 |
| TSA (1h) | 1715 | 2.2E+07 | 1.3E+04 |
| **Total** | **11848** | **1.6E+08** | |

**Supplementary Table 4: Number HeLa cells removed during data cleanup across all conditions**

| | n | % total |
|---|---|---|
| **All (non-border) cells imaged** | **3161** | **100%** |
| Mitotic cells (excluded) | 193 | 6.1% |
| Polynucleated cells (excluded) | 80 | 2.5% |
| Extreme DNA content (excluded) | 101 | 3.2% |
| Acquisition error (excluded) | 43 | 1.4% |
| Background artefacts (excluded) | 10 | 0.3% |
| **Final analysed cells** | **2731** | **86.4%** |

**Supplementary Table 5: Number of HeLa cells and pixels analysed per condition**

| Condition | Number of cells | Number of pixels (cell) | Mean area (cell) |
|---|---|---|---|
| Scrambled siRNA | 2301 | 1.8E+08 | 7.8E+04 |
| SBF2 siRNA | 430 | 9.1E+07 | 2.1E+05 |
| **Total** | **2731** | **2.7E+08** | |

**Supplementary Table 6: Number of cells and molecular profiles per split used for dataset training.**

| | train | val | test |
|---|---|---|---|
| **184A1 dataset** | | | |
| cells | 9’472 | 1’181 | 1’195 |
| subsampled molecular profiles | 615’240 | 76’432 | 76’956 |
| **HeLa dataset** | | | |
| cells | 2’160 | 284 | 287 |
| subsampled molecular profiles | 10’628’809 | 1’394’437 | 1’390’160 |

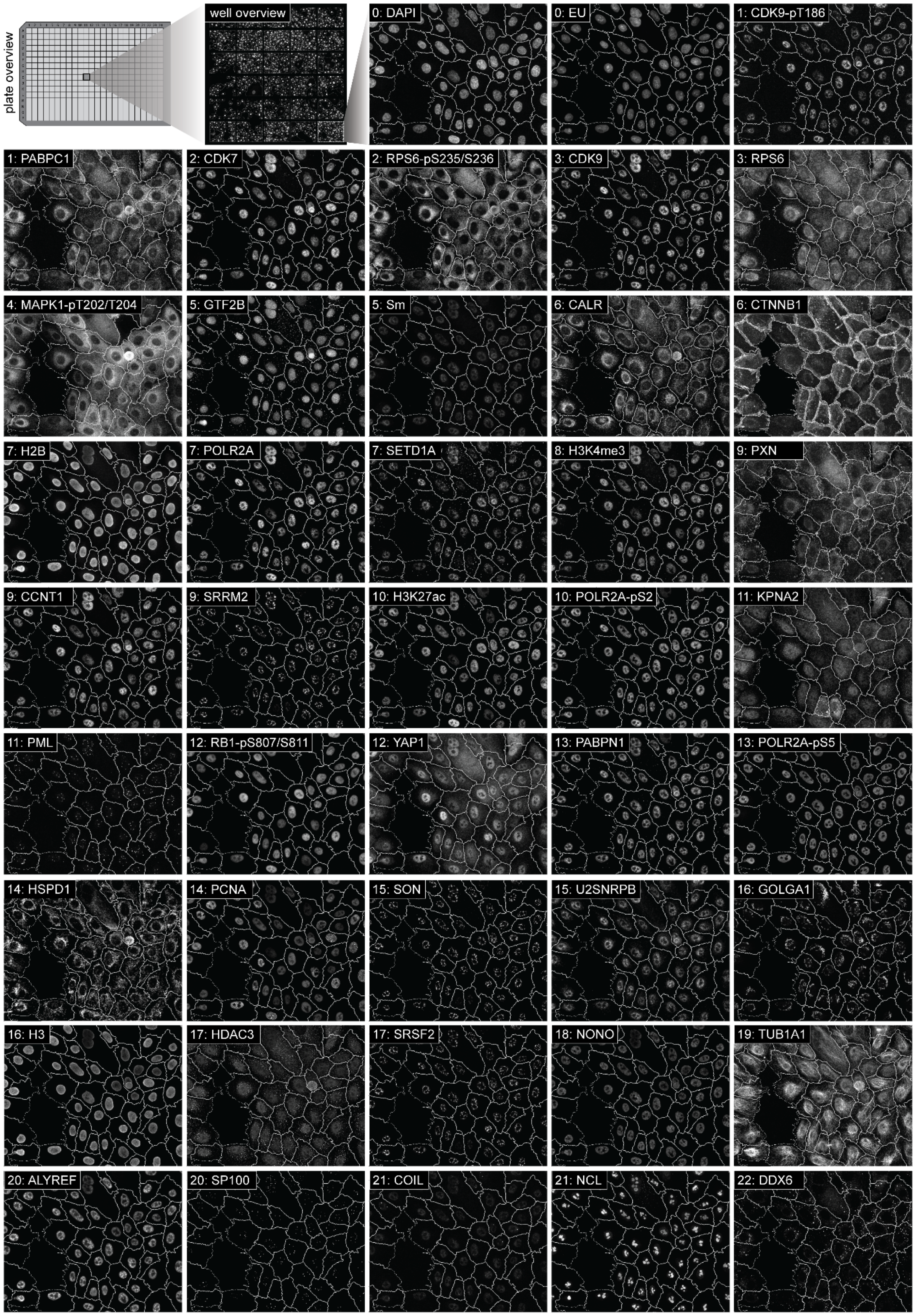


**Supplementary Figure 1: Iterative Indirect Immunofluorescence Imaging (4i) at high spatial resolution.** Example images with overlaid cell segmentation of 184A1 cells for each of the 43 channels measured in by 4i.

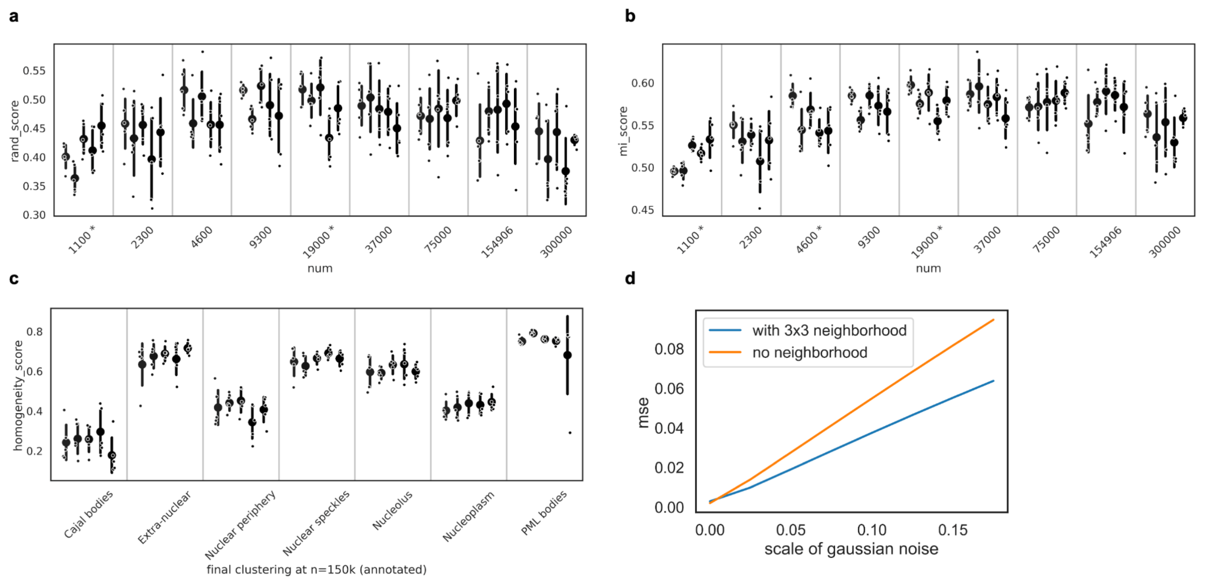


**Supplementary Figure 2: Robustness to noise and randomness in subsampling and clustering using the 184A1 dataset.**

**a,b**: Cluster stability at different subsample sizes. For each subsample size clusterings from 5 different subsamples and 5 different Leiden initialisations in each subsample are computed and compared to the final CSLs obtained at 150k samples. Cluster overlap is assessed with adjusted rand index (a) and adjusted mutual information (b). Scores are grouped by subsample with 95% confidence intervals obtained by bootstrapping. An asterisk * at a subsample size indicates a significant ($p<0.05$) effect of the subsample on the score, obtained by ANOVA. Different subsamples have no significant influence on cluster stability for subsamples sizes >= 37k.

**c**: Homogeneity of different clusterings at 150k samples w.r.t. to final CSLs. Homogeneity is high for a certain CSL, if for each cluster in a clustering, all points assigned to this cluster either belong or not belong to the CSL. Homogeneity was lowest for Nucleoplasm and Nuclear periphery, and Cajal bodies, showing that the boundaries between these clusters are not as clearly defined from the data as for CSLs Nuclear Speckles, PML bodies, Nucleolus.

**d**: Robustness of cVAE latent space when adding Gaussian noise to the input. Scale is the standard deviation of the Gaussian distribution. Shown is the MSE and 95% confidence interval of pairwise comparisons with 5 different random seeds. cVAE latent spaces obtained by training with a local 3x3 neighbourhood as context for every molecular profile and without any neighbourhood information are compared. Using neighbourhood information adds to increased stability of the latent space.

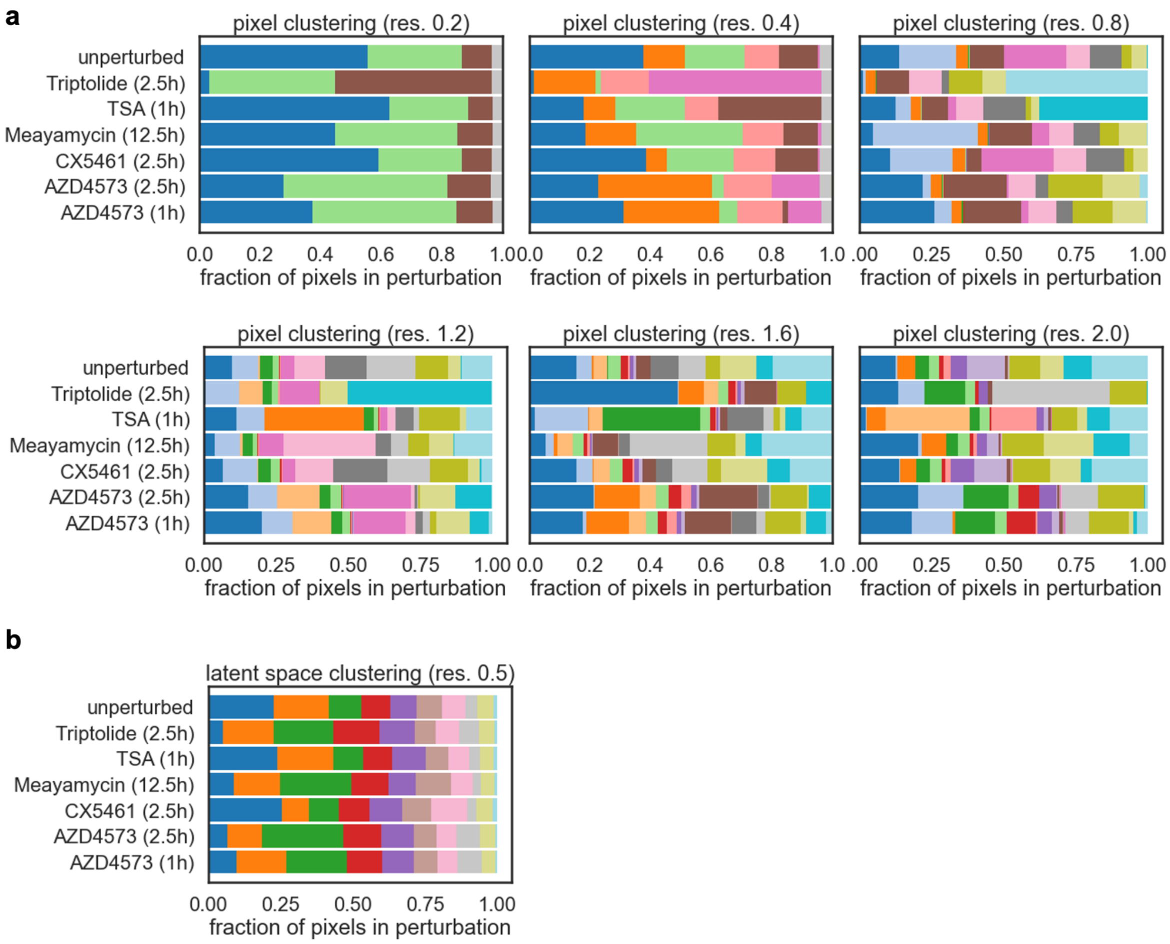


**Supplementary Figure 3: Pixel clustering is perturbation-dependent at different leiden clustering resolutions.**

**a:** Fraction of pixels assigned to each cluster per perturbation colored by direct pixel clustering for leiden resolutions 0.2,0.4,0.6,0.8,1.2,1.6,2.0

**b:** Fraction of pixels assigned to each cluster per perturbation colored by latent space clustering before annotation.

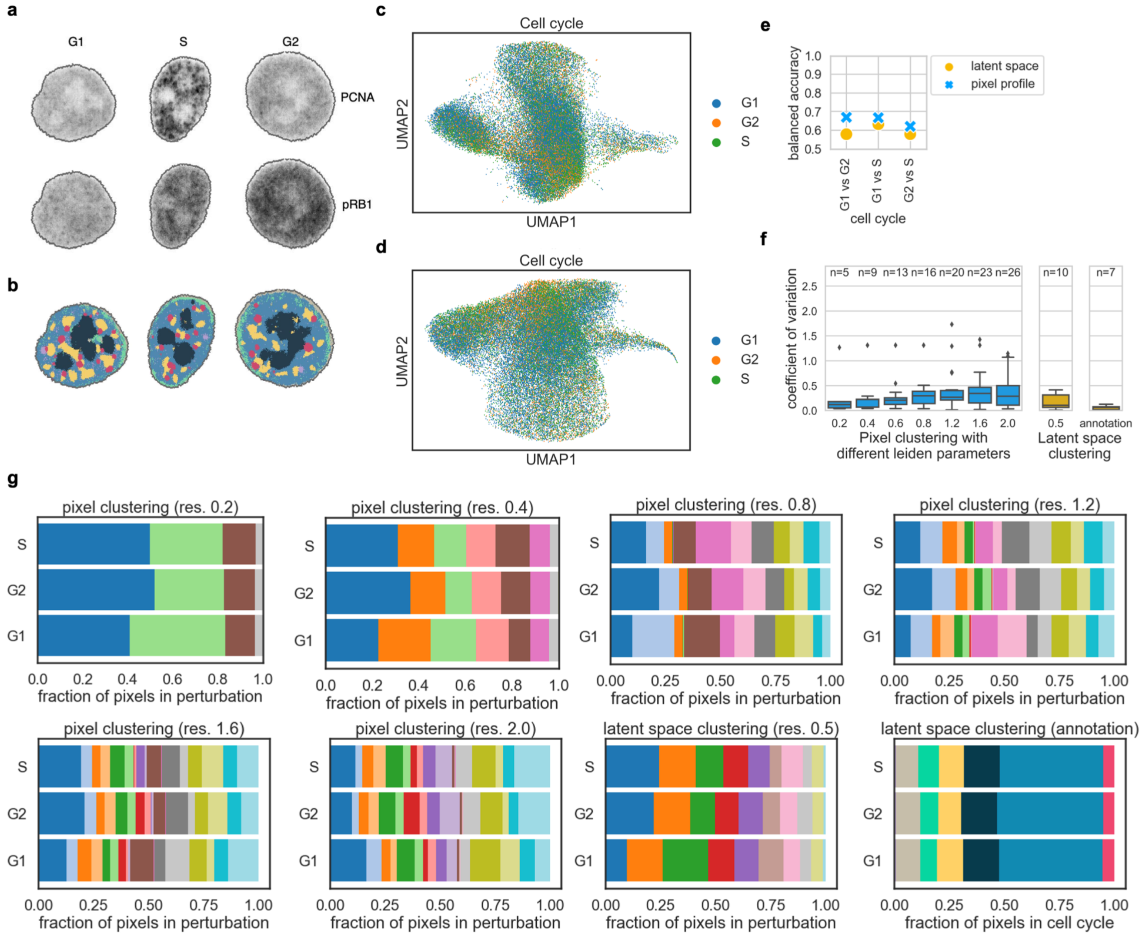


**Supplementary Figure 4: Pixel clustering is cell-cycle-dependent at different leiden clustering resolutions.**

**a**: Example cells from G1, S, and G2 phase colored by cell-cycle specific markers PCNA and pRB1.
**b**: Same cells colored by CSL clusters.
**c-d**: UMAP representation of multiplexed pixel profiles from unperturbed cells (c) and of corresponding cVAE latent space (d) colored by cell cycle.
**e:** Comparison of cell-cycle-specificity of 4i pixel profiles and cVAE latent space coordinates. Plots show balanced accuracy scores of pairwise binary logistic regression classifiers predicting cell cycle from normalised 4i pixel profiles or latent representations of pixels. Accuracy values of 0.5 indicate random chance (perturbation information is not present in the data). Latent space contains less cell-cycle information than pixel profiles.
**f**: Comparison of cell-cycle-specificity of direct clustering at different leiden resolutions (0.2, 0.4, 0.6, 0.8, 1.2, 1.6, 2.0) with cVAE latent space clustering. Plots show coefficient of variation of the fraction of pixels in each cell-cycle stage assigned to each cluster. Boxplot summarises results for all clusters with resulting cluster number n shown above.
**g:** Fraction of pixels assigned to each cluster per cell-cycle stage colored by pixel profile clustering for resolutions 0.2,0.4,0.6,0.8,1.2,1.6,2.0 and by latent space clustering and annotation.

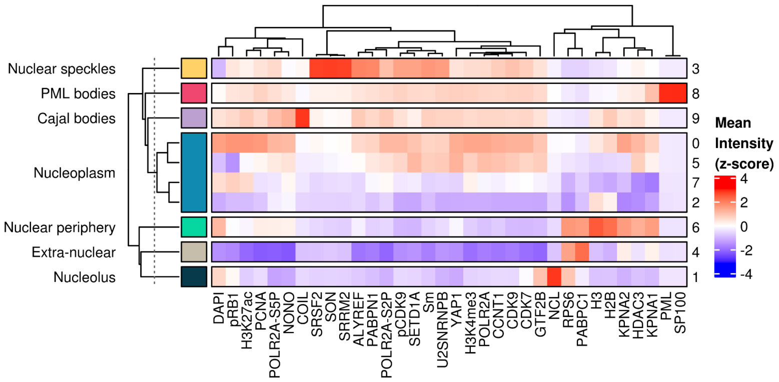


**Supplementary Figure 5: Mean intensities of each channel across different clusters (unperturbed 184A1 cells).**

Original Leiden clusters obtained from cVAE latent representation are shown on the right and their corresponding annotation is shown on the left. Four original clusters are manually merged into the "Nucleoplasm" CSL. Values z-scored by channel (cf Fig 2b).

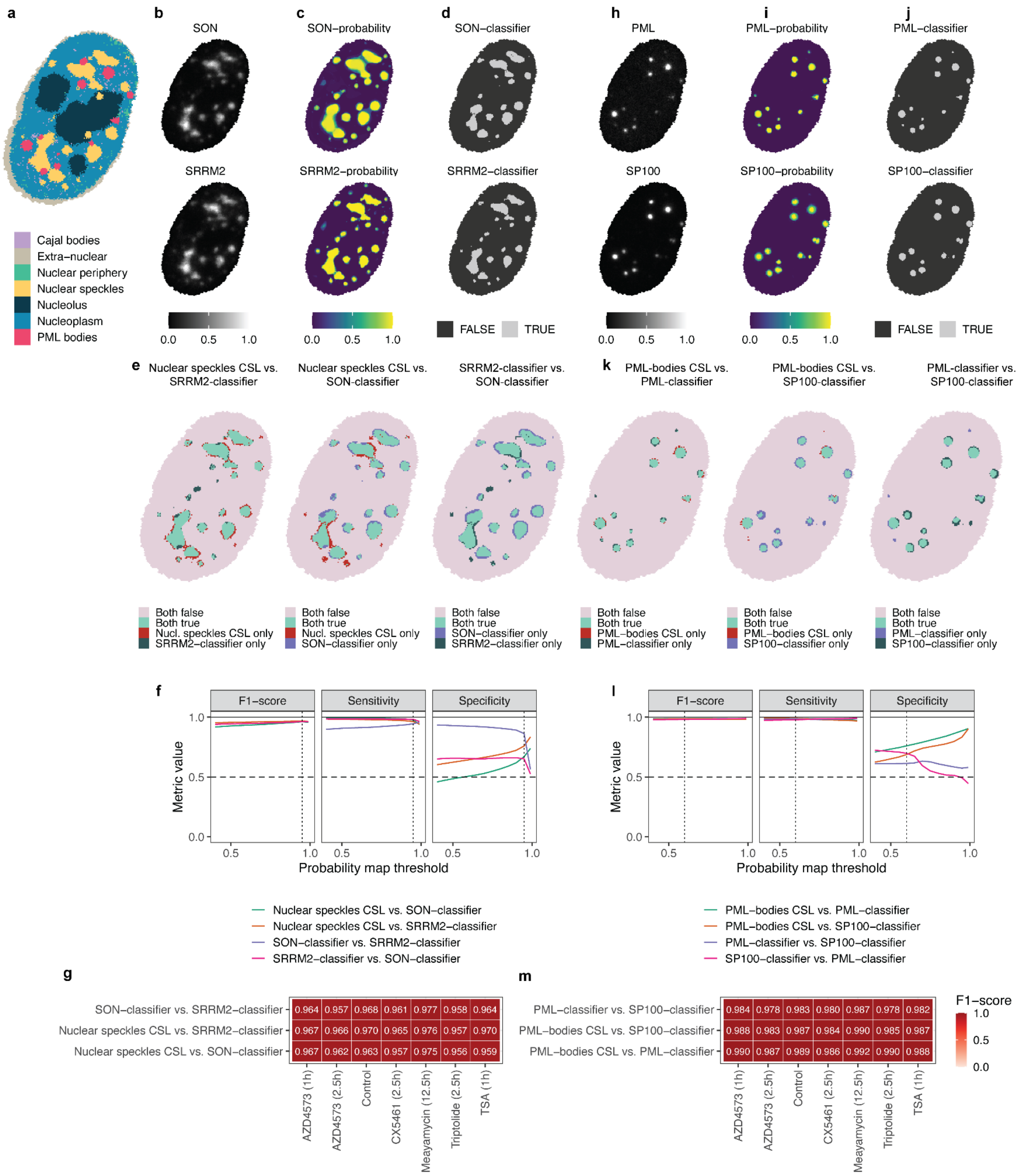


**Supplementary Figure 6: Comparison of CSL-derived nuclear speckles/PML bodies with supervised segmentation of nuclear speckles/PML bodies.**

**a:** CSLs generated using CAMPA for an example 184A1 nucleus.

**b:** Measured intensities of canonical nuclear speckle markers SON and SRRM2.

**c:** Probability maps of nuclear speckles generated through supervised pixel clustering of single-channel images in Ilastik.

**d:** Nuclear speckle classifier derived from thresholding the probability map at P=0.95.

**e:** Comparison of pixels assigned to nuclear speckles using supervised classifiers based on SON, SRRM2 with nuclear speckle CSL.

**f:** Quantitative comparison of the different classification approaches using SON and SRRM2 classifiers as the ground-truth. x-axis shows the value at which the probability map is thresholded to generate the segmentation. Dashed line at P=0.95 is the classification shown in D. Metrics computed on a random 10% subsample of all data.

**g:** $F_1$-scores for each comparison for all perturbations individually. Data from all pixels in a single well for each condition. Probability map threshold at P=0.95.

**h-j:** As in b-d for canonical PML body markers SP100 and PML. Classification in i is the probability map thresholded at P=0.6.

**k:** As in e for PML bodies.

**l:** As in f for PML bodies. Dashed line at P=0.6 is the classification shown in j.

**m:** As in g for PML bodies. Probability maps thresholded at P=0.6.

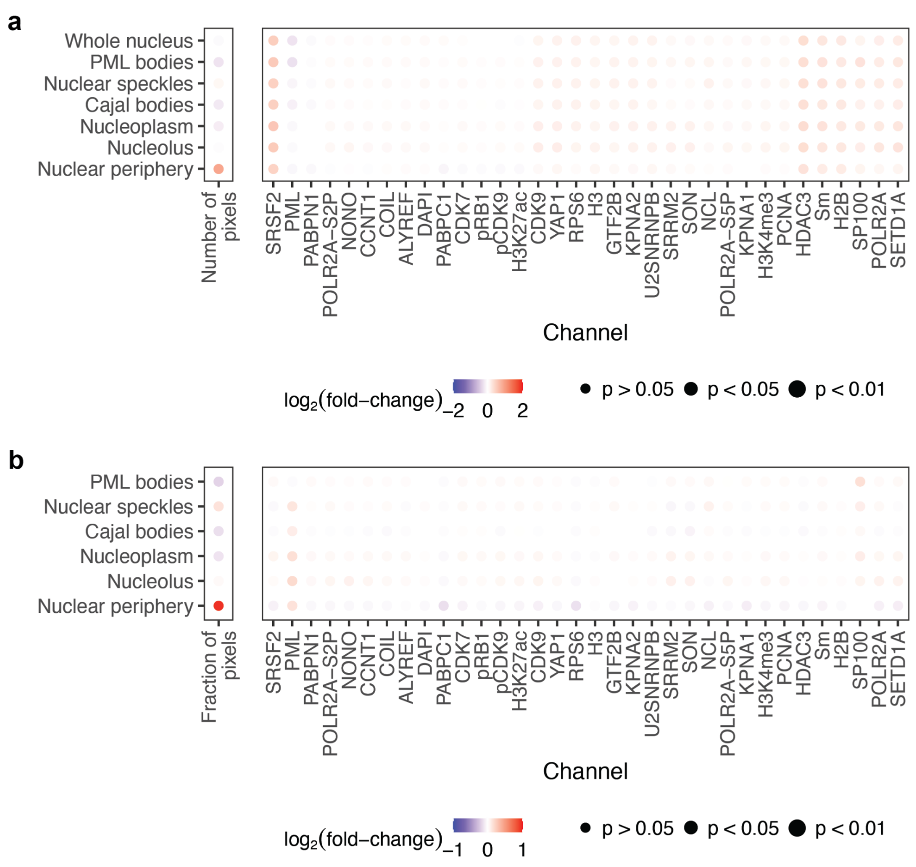


**Supplementary Figure 7: Mean intensity changes upon DMSO treatment.**

**a**: Log2 fold-change of mean intensities for each channel in each CSL, or number of pixels in each CSL, when comparing DMSO with untreated cells. P-values show significance of DMSO treatment on protein levels for each channel/CSL, as determined from the mixed effect model. P-values are corrected for multiple hypothesis testing using the Benjamini-Yuketeli method. DMSO and untreated cells show similar mean intensities across all CSLs and are pooled as ‘unperturbed’ cells in subsequent analyses.

**b**: As above, except normalised by the overall (whole-nucleus) changes in intensity. In this case p-values indicate significance of mean intensity change in CSL compared to the change observed for the whole nucleus (see Methods).

a

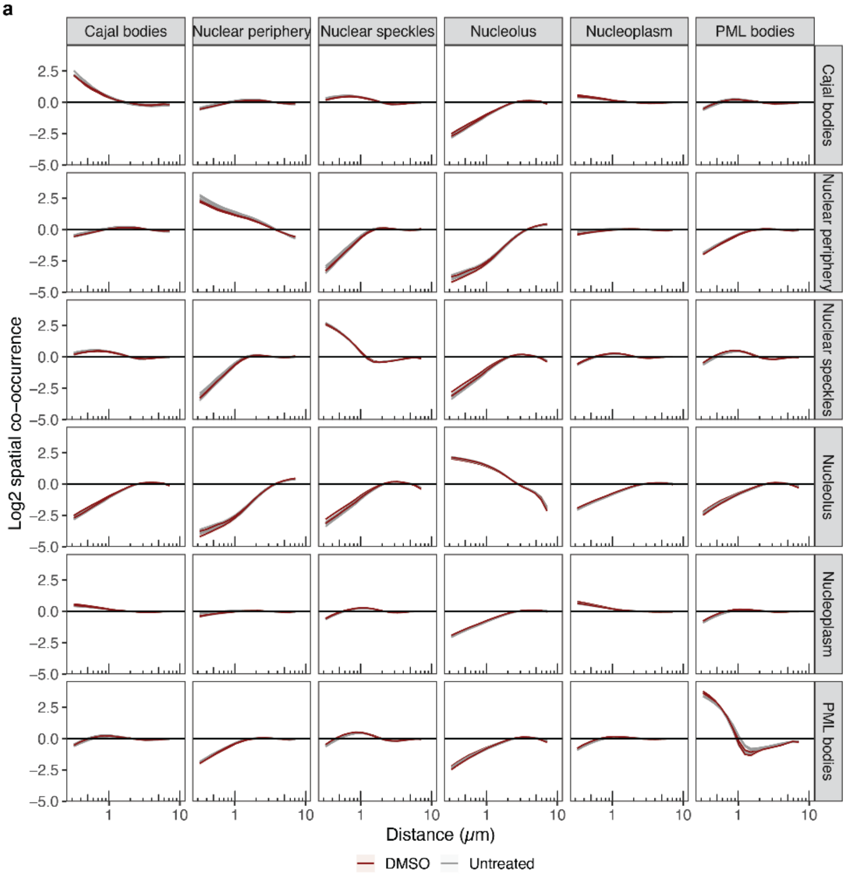

Cajal bodies
Nuclear periphery
Nuclear speckles
Nucleolus
Nucleoplasm
PML bodies
Log2 spatial co-occurrence
2.5
0.0
−2.5
−5.0
1
10
Distance (μm)
DMSO
Untreated


b

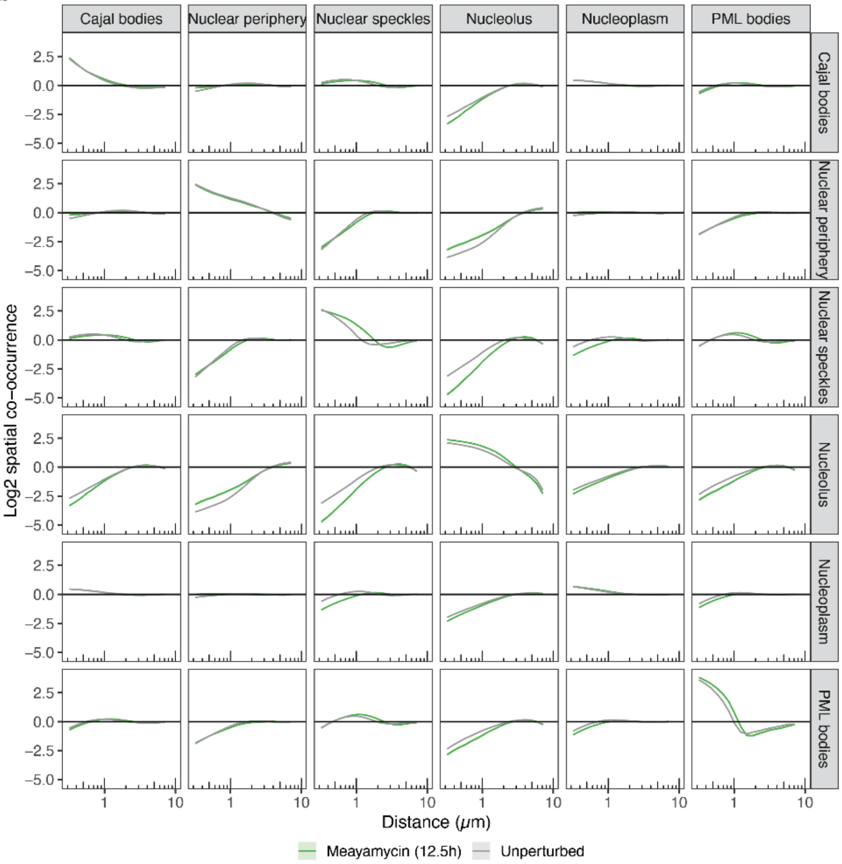

Cajal bodies
Nuclear periphery
Nuclear speckles
Nucleolus
Nucleoplasm
PML bodies
Log2 spatial co-occurrence
2.5
0.0
−2.5
−5.0
1
10
Distance (μm)
Meayamycin (12.5h)
Unperturbed

**Supplementary Figure 8: Pairwise co-occurrence scores between CSLs.**

**a**: Pairwise mean log2 spatial co-occurrence between all CSLs as a function of distance on x-axis (on log scale) for DMSO and untreated control wells (pooled as 'Unperturbed' in Fig 3h). Each well plotted as an individual line. Shaded regions indicate 95% confidence interval for the mean. DMSO and untreated cells show similar co-occurrence scores and are pooled as 'unperturbed' cells in subsequent analyses.

**b**: Pairwise mean log2 spatial co-occurrence between all CSLs as a function of distance on x-axis (on log scale) for Meayamycin-treated and unperturbed cells. Shaded regions indicate 95% confidence intervals for the mean.

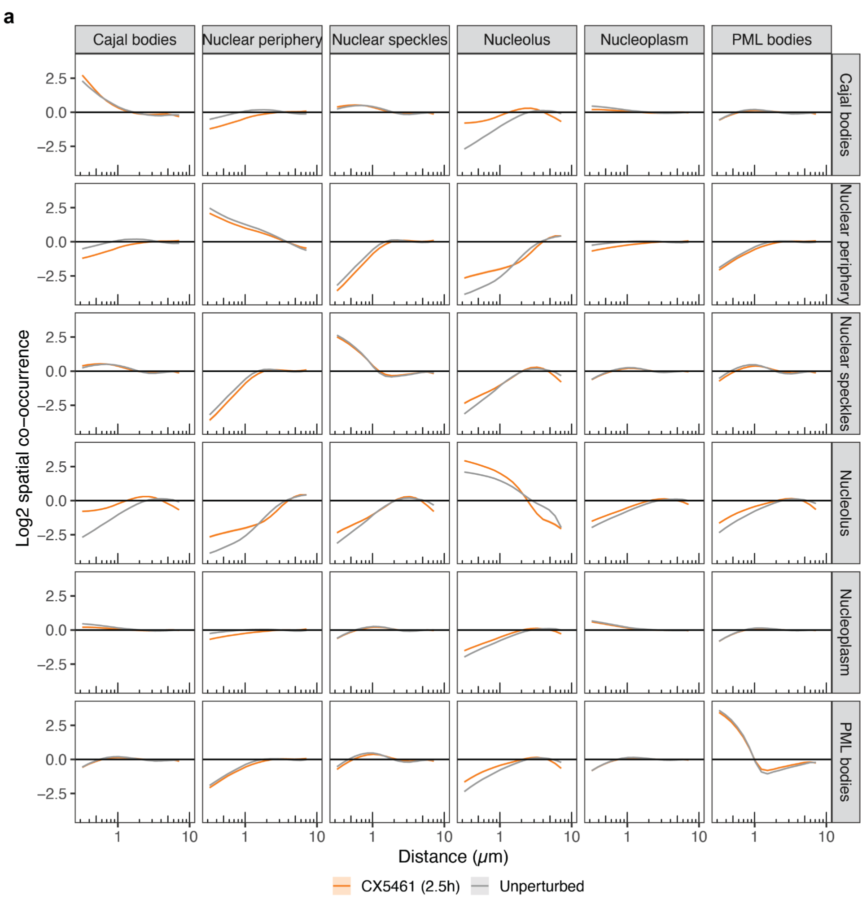


**Supplementary Figure 9: Pairwise co-occurrence scores between CSLs.**
**a**: Pairwise mean log2 spatial co-occurrence between all CSLs as a function of distance on x-axis (on log scale) for CX5461-treated and unperturbed cells. Shaded regions indicate 95% confidence intervals for the mean.

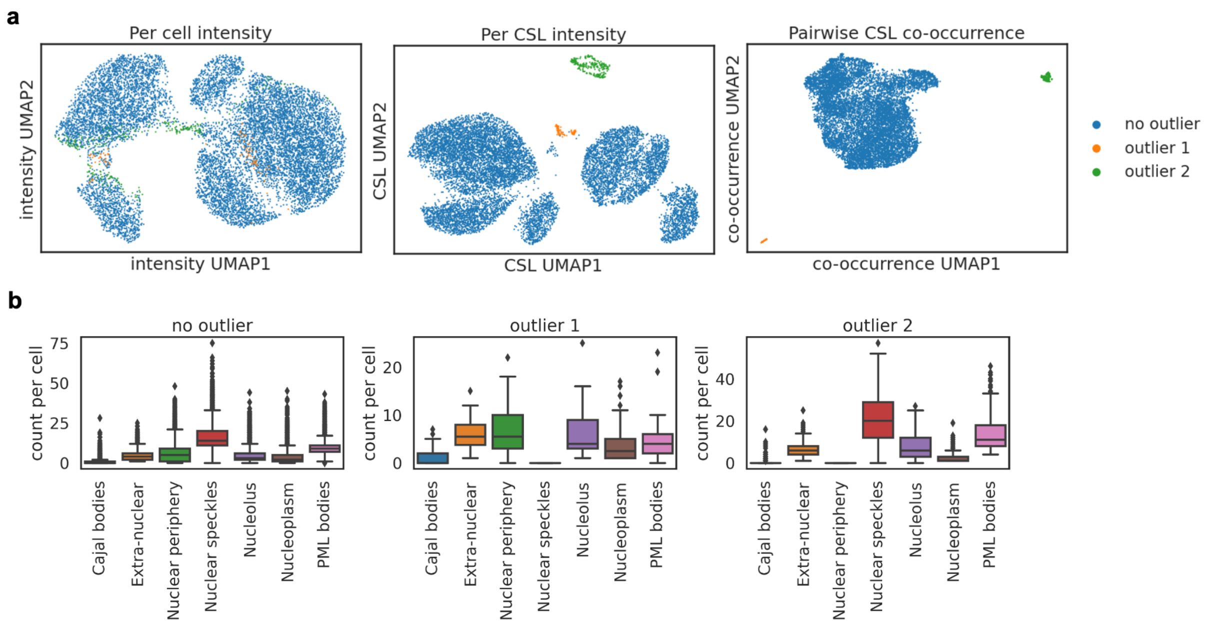


**Supplementary Figure 10: Examination of outlier cells excluded in Fig. 4 on the basis of the co-occurrence UMAP.**

Two outlier clusters were identified (outlier 1: 88 cells, outlier 2: 256 cells).

**a:** UMAP embedding of cells using different cellular features. Points colored by outlier clusters. The identified outlier clusters are visible outliers on the intensity UMAP and the co-occurrence UMAP.

**b:** Barplot of number of individual CSLs per cell, colored by CSL, for non-outlier cells, and the two outlier clusters. Cells in outlier cluster 1 do not contain the Nuclear speckle CSL, and cells in outlier cluster 2 do not contain the Nuclear periphery CSL.

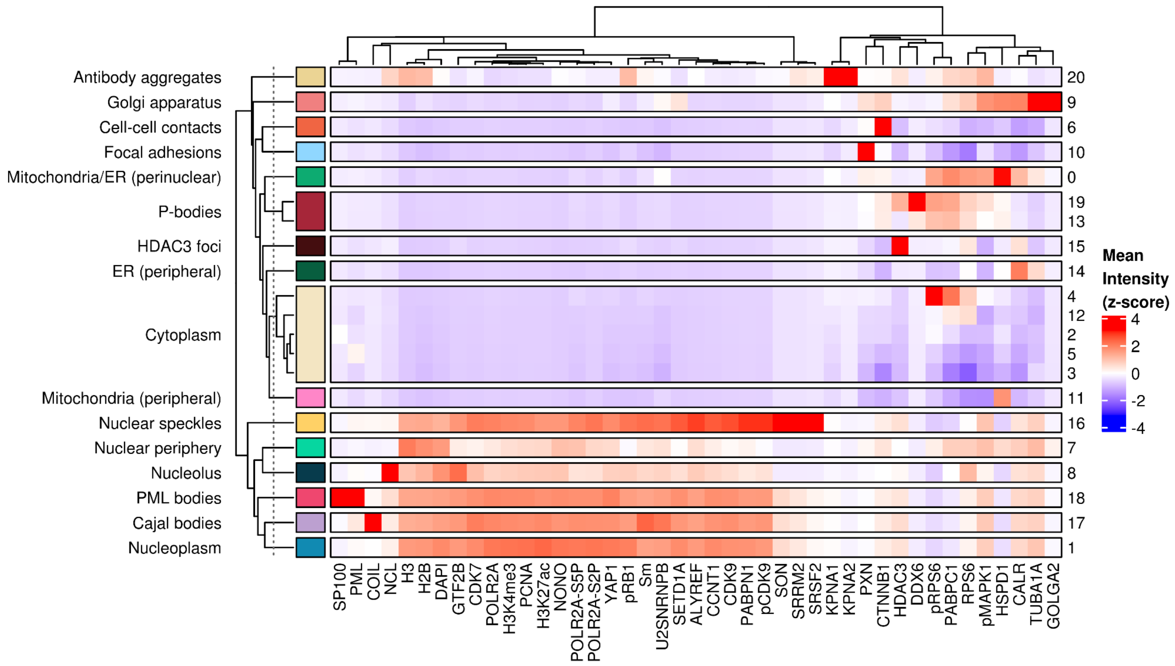


**Supplementary Figure 11: Mean intensities of each channel across different clusters (HeLa cells, scrambled siRNA).**

Original Leiden clusters obtained from cVAE latent representation are shown on the right and their corresponding annotation is shown on the left. Five original clusters are manually merged into the “Cytoplasm” CSL. Two original clusters are manually merged into the “P-bodies” cluster. Values z-scored by channel (c.f. Fig 5b).

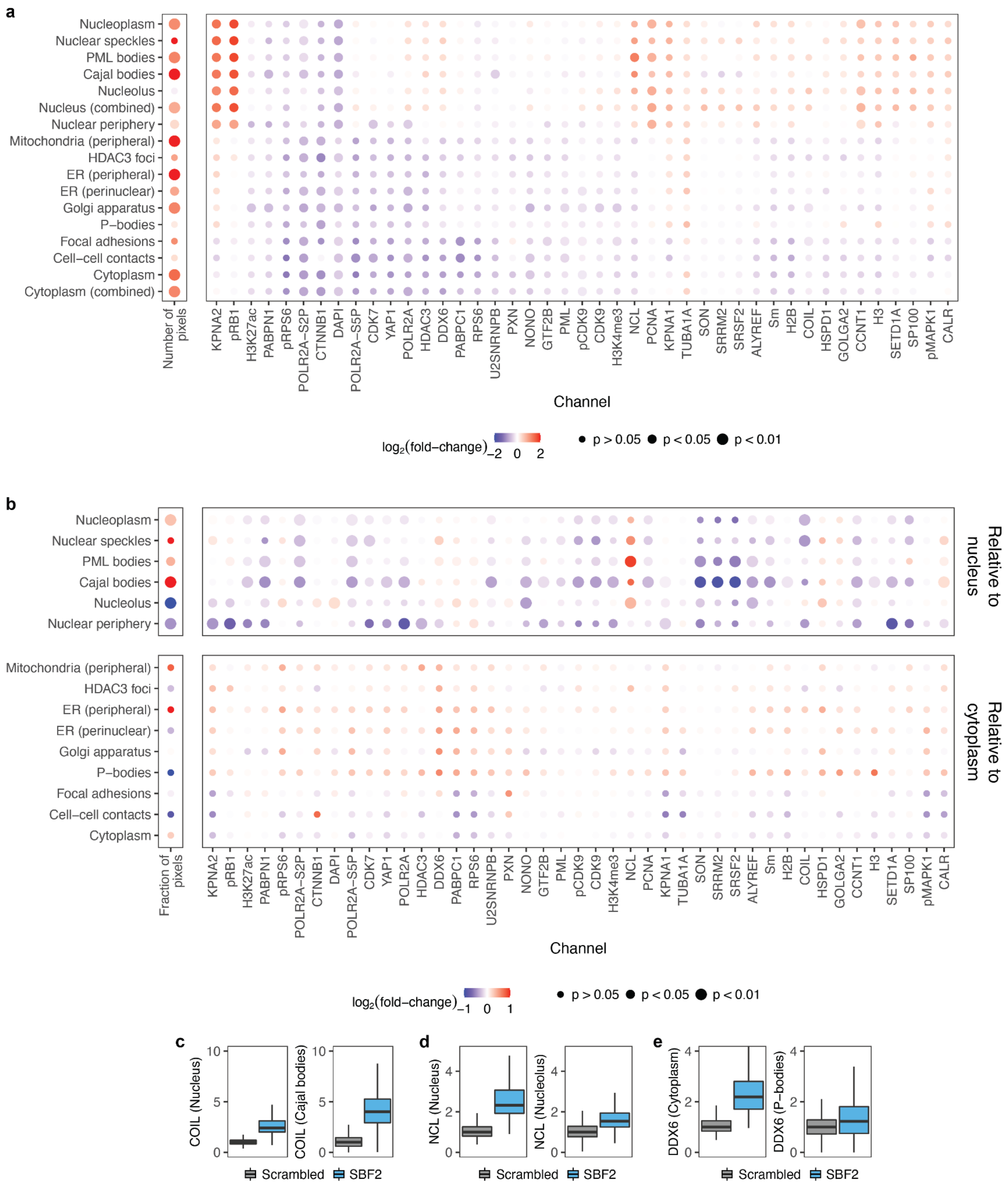


**Supplementary Figure 12: Quantification of mean intensity changes of each channel in each CSL in HeLa cells upon SBF2-knockdown**

**a**: Log2 fold-change of mean intensities for each channel in each CSL, or number of pixels in each CSL, when comparing *SBF2*-knockdown with scrambled siRNA control cells. P-values show significance of *SBF2*-knockdown on protein levels for each channel/CSL, as determined from the mixed effect model. P-values are corrected for multiple hypothesis testing using the Benjamini-Yuketeli method.

**b**: As above, except normalised by the whole-nucleus or whole cytoplasm changes in intensity, respectively for the upper and lower panels. In this case p-values indicate significance of mean intensity change in CSL compared to the change observed for the nucleus or cytoplasm, respectively (see Methods).

**c**: Sum intensity of COIL in the nucleus or Cajal bodies of scrambled and SBF2-knockdown HeLa cells. Boxplots summarise distributions across cells.

**d**: As in c for NCL/Nucleolus.

**e**: As in c for DDX6/P-bodies.

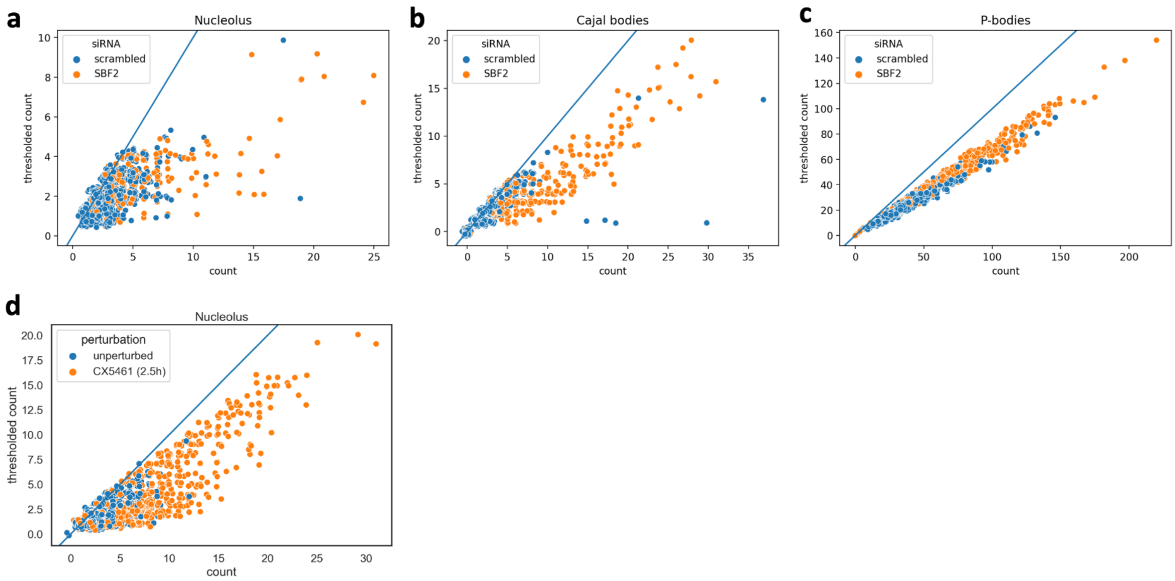


**Supplementary Figure 13: Effect of size-filtering on CSL counts per cell.**
Number of CSLs per cell before and after filtering small individual CSLs for different CSLs (<10px or cumulative sum of removed object <10% of total CSL area in this cell). From cells that contain more individual CSLs before filtering, more individual CSLs are removed, indicating that these cells contained many small individual CSLs. Blue line indicates diagonal. Small jitter (sigma=0.2) applied to each point for visualisation.

**a:** Nucleolus count in scrambled and SBF2-knockdown HeLa cells.

**b:** Cajal body count in scrambled and SBF2-knockdown HeLa cells.

**c:** P-body count in scrambled and SBF2-knockdown HeLa cells.

**d:** Nucleolus count in unperturbed and CX5461 184A1 cells.

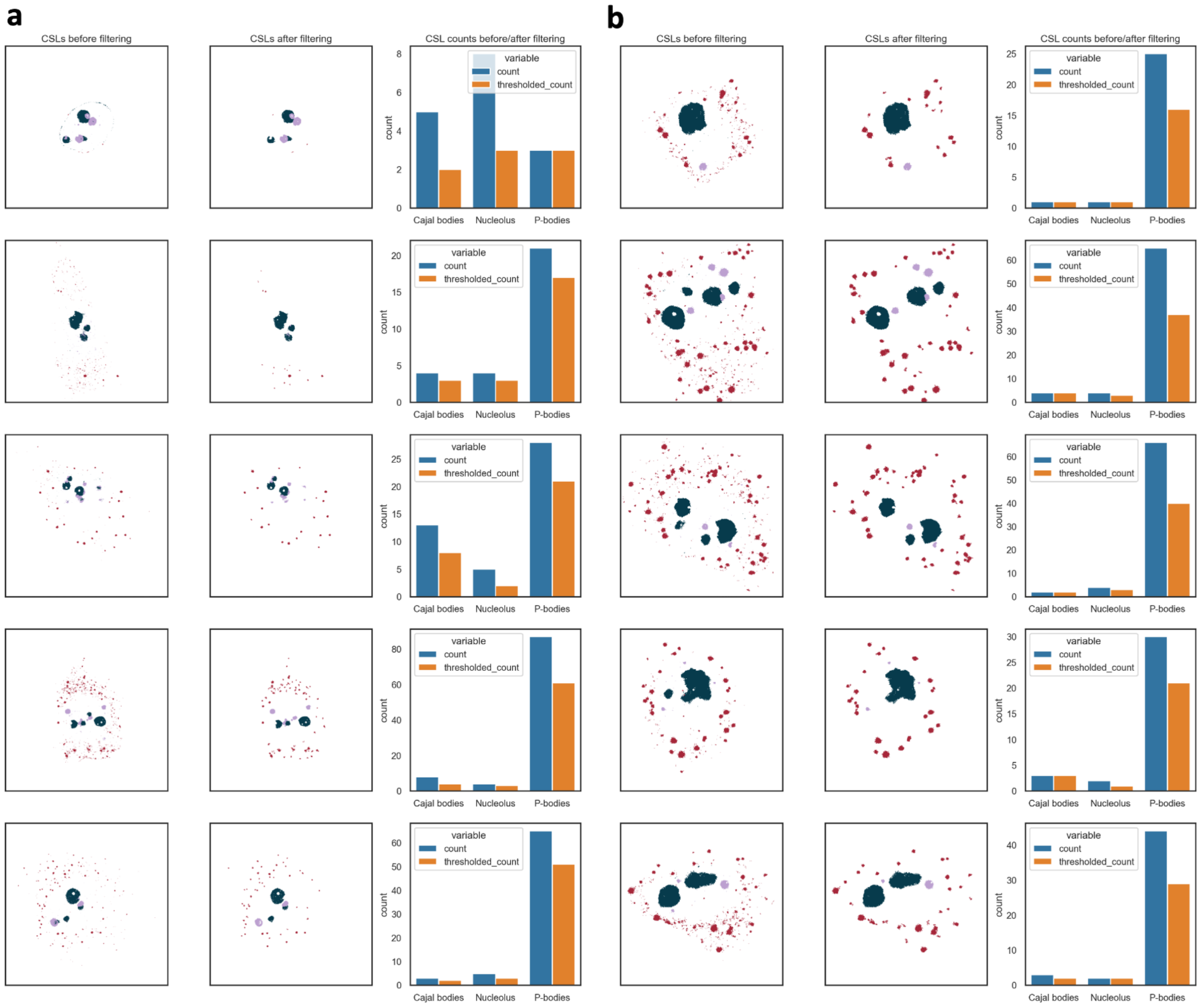


**Supplementary Figure 14: Example cells showing CSLs before and after size-filtering.** Cajal bodies, Nucleolus and P-bodies and their counts before and after filtering (which removes bodies <10px or those whose cumulative sum is <10% of total CSL area per cell) in HeLa cells (see Methods). Note that plot sizes differ between a and b (a: 700x700px, b: 400x400px).

**a:** Randomly selected cells from SBF2-knockdown cells. Left: Nucleolus (dark blue), Cajal body (purple), and P-body (red) CSLs before size-filtering. Middle: CSLs after size-filtering. Right: Barplot of CSL counts before and after filtering.

**b:** Same as in panel a, for scrambled cells but for randomly selected cells from scrambled condition.

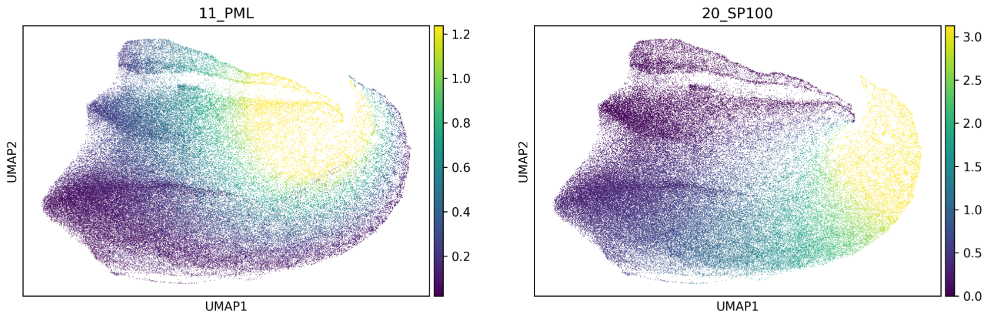


**Supplementary Figure 15: Heterogeneity of PML bodies.**
UMAP of cVAE latent representation of 10% of all PML-body pixels from scrambled HeLa cells. Pixels are colored by PML-body markers SP100 (left) and PML (right). PML-body pixels are heterogeneous, from PML-enriched to SP100-enriched, and the latent representation conserves this heterogeneity.
